# A Computational Re-evaluation of Spatial Trials for Zoonotic Tuberculosis Control: Model Misspecification, Diagnostic Misclassification, and the Illusion of Wildlife Culling Efficacy

**DOI:** 10.64898/2026.08.29.747978

**Authors:** Paul R Torgerson

## Abstract

1. Wildlife reservoir management frequently relies on the Randomised Badger Culling Trial’s (RBCT) trade-off hypothesis, which posits that reductions in cattle herd infections are offset by a perturbation effect driven by disrupted host dispersal. This paper evaluates the computational and epidemiological robustness of this historical trial, which serves as the foundational empirical experiment guiding zoonotic tuberculosis (*Mycobacterium bovis*) control policies.
2. Using generalized linear mixed models with a generalized Poisson error distribution to explicitly address historical data overdispersion, this study contrasts traditional parametric inference against exact cluster-constrained permutation tests across distinct operational definitions of disease incidence.
3. Non-parametric diagnostics reveal that previously reported treatment and perturbation effects render as statistical artifacts under exact non-parametric permutation. Inside culling zones, parametric significance fails to withstand exact permutation verification due to extreme data leverage in localized cluster blocks.
4. Crucially, when diagnostic misclassification biases are eliminated by analysing total reactor datasets, all apparent culling effects disappear, and information criteria overwhelmingly favour nested null architectures. Unconfirmed reactors likely represent true biological infections missed by low-sensitivity post-mortem macro-necropsy, proving that host removal tracks observation noise rather than genuine zoonotic transmission pathways.
5. Finally, empirical scaling conducted in this study identifies a novel mathematical saturation effect, demonstrating that this sub-linear scaling is an operational artifact of unmodelled herd-level disease recurrence over time.
6. **Policy implications.** Because current zoonotic tuberculosis intervention frameworks are built upon a structurally misspecified statistical model, they have driven large-scale veterinary policies resulting in substantial, unevidenced ecological and economic interventions while failing to provide genuine public health, animal health, or disease control benefits.

## 1. Introduction

The management of endemic wildlife reservoirs of zoonotic pathogens represents one of the most contentious interfaces of public health, veterinary policy, and agricultural economics. A primary example is the control of bovine tuberculosis (bTB), caused by *Mycobacterium bovis,* where the role of the European badger (*Meles meles*) as a potential maintenance host has driven decades of controversial badger culling policy in the British Isles.

The design of effective public policy for infectious disease control hinges on the integrity of the diagnostic metrics used to define a case. In the epidemiology of bTB, the standard unit of regulatory interest is the incidence of herd breakdowns. The Randomised Badger Control Trial (RBCT) was an expensive (£49 million) and wide-scale intervention experiment with the aim at quantifying the effect of badger removal on the herd incidence of tuberculosis in cattle herds in England. The results of the RBCT were published in *Nature* in 2006 (Donnelly et al., 2006), with a claim that there was a strong positive effect of badger culling. That is proactive culling of badgers reduced the incidence of herd breakdowns by approximately 20%. There was also a claim that there was a negative effect that resulted in an increase in herd breakdowns in neighbouring areas caused by infected badgers migrating away from areas where culling was taking place (the “perturbation” effect”). This evidence led to an expensive and ethically questionable programme of badger culling which commenced in 2013 as part of the control programme for bTB. Since then over 250,000 badgers have been culled at great expense using free shooting methods that can cause slow and painful death. Questions in parliament and a recent parliamentary debate have occurred as a result of the badger culling programme. The latest bTB strategy review (Cross et al., 2026) recommendations to the UK Department for Environment, Food and Rural Affairs (DEFRA) - a ministerial department of the United Kingdom government responsible for environmental protection, food production and standards, agriculture, fisheries, and rural communities, primarily within England. This report continues to cite the RBCT as evidence for the efficacy of badger intervention for control of bTB.

Details of the RBCT experiment are available elsewhere (Donnelly et al., 2006; Bourne et al., 2007; Donnelly et al., 2007). Briefly there were 10 ‘triplets’. Each triplet had three areas of approximately 100 km^2^. In each triplet, there was a proactive cull area where the aim was to pre-emptively remove most of the badgers, a reactive cull area where badgers were removed in the vicinity of a herd breakdown, and a control area where badgers remained undisturbed. The reactive arm of the trial was abandoned before completion (Donnelly et al., 2003).

Evaluation of the experiment partitioned breakdowns into two administrative classifications: “confirmed” and “unconfirmed” incidents. In the RBCT, a breakdown is designated as confirmed if at least one animal within a herd presents visible tuberculosis lesions during post-mortem inspection or yields positive bacteriological cultures. Conversely, an unconfirmed breakdown denotes a herd where at least one animal reacts positive to the primary live-animal diagnostic but fails to present macroscopic or culture positive lesions at slaughter (Bourne et al., 2007).

This binary partition underpins a major vulnerability in historical trial interpretations. From a veterinary standpoint, the routine Single Cervical Comparative Intradermal Tuberculin Test (SICCT) operates with an exceptional field diagnostic specificity of close to 100% (Goodchild et al., 2015; Nuñez-Garcia et al., 2018). Because the probability of a false-positive reactor is very low, such reactors are nearly entirely true biological hosts of *M bovis.* The systemic divergence between skin-test positivity and post-mortem confirmation is not driven by live-animal diagnostic failure, but rather by the severe diagnostic limitations of standard slaughterhouse macro-necropsy. It has been established that post-mortem examination has an imperfect diagnostic sensitivity ranging between 46% for routine slaughterhouse surveillance to 79% for detailed postmortem examination (Nuñez-Garcia et al., 2018). It can also be highly variable between abattoirs (Pascual-Linaza et al., 2017).

In the structural specification of macro-epidemiological models for livestock diseases, modelling the relationship between host population density and transmission opportunity is a central challenge. Standard epidemiological formulations traditionally rely on the assumption that the rate of new infections scales proportionately with exposure. In mixed-effects regression frameworks, this linear proportionality assumption is imposed by assigning log-transformed exposure metrics—such as herd density and/or total time-at-risk—as a fixed statistical offset (*offset* (log (*N ×T*)) This restricts the associated regression coefficient to a fixed value of unity (*β* = 1).

While mathematically parsimonious, treating exposure as a rigid fixed offset assumes that doubling the population size or the observation window doubles the biological opportunities for transmission. In complex multi-host epidemics, such as hypothesised *M. bovis* transmission at the wildlife-livestock interface, this linear scaling property may vary. To address this limitation, contemporary spatial epidemiologists can treat exposure as a free log-linear explanatory variable *β* log (*N ×T*), allowing the data to determine the scaling exponent empirically. If the resulting regression coefficient is above unity (*β* >1), then it is evidence that transmission rate disproportionately increases as population density and/or time at risk increase (ie a density dependence). In contrast, when the resulting regression coefficient falls significantly below unity (β < 1), it provides evidence of a transmission saturation effect.

## 2. Materials and Methods

### 2.1 Data and analysis

All data were taken from the supplementary material of Donnelly et al. (2006). It is reproduced in supplementary data file 1. All analysis was undertaken in R (R Core Team, 2026). The statistical code is provided as additional material (supplementary file 2).

#### 2.1.1 The Historical Fixed-Effects Poisson Baseline (Donnelly et al. 2006)

This approach is structured as a classic parametric fixed-effects Poisson generalized linear model (glm):

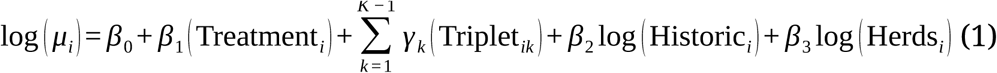

This framework enforces a strict Poisson mean-variance identity (φ = 1) and entirely omits temporal exposure profiles (time at risk) despite highly variable observation windows across the randomized trial blocks. Model misspecification, residual scaling anomalies, and quantile deviations for this fixed-effects baseline were formalized using simulated quantile residuals via the DHARMa package (Hartig, 2024).

### 2.2 Generalized Mixed Model Specification

To evaluate the impact of wildlife intervention (badger culling) on bTB herd breakdowns, a generalized linear mixed model (GLMM) was formulated using a generalized Poisson error distribution to account for over-or under-dispersion within count data. The baseline observed full model is parametrised as:

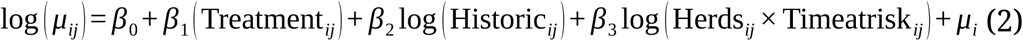

where *μ*_(_*_ij_*_)_ represents the expected bTB incidence for herd j within the i-th randomized cluster block (triplet). The term 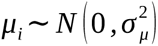 represents a random intercept for triplet to control for unobserved regional spatial and temporal clustering. The model incorporates continuous historical baseline risks log (*Historic*) and log-transformed exposure log (*Herds ×Timeatrisk*). This was implemented in R using the glmmTMB package (Brooks et al., 2017).

Models were fitted using two distinct dependent variables to evaluate the sensitivity of the policy metrics:

1. **Confirmed breakdowns only:** Incidents in which postmortem examination of slaughtered cattle led to detection of TB lesions or culture of *M. bovis* in at least one animal in the herd.
2. **Total Breakdown Dataset:** Herd incidents combining both confirmed and unconfirmed breakdowns. An unconfirmed breakdown is where at least one animal in the herd has a positive result from the routine surveillance skin test (a “reactor”), but the infection was not confirmed at post mortem.

### 3. Statistical Estimation of Potential False Positive Herds

Routine testing of cattle herds was conducted using the single intradermal comparative cervical tuberculin test (SICCT or ‘tuberculin skin test’). The operation and interpretation of the test is described in full in Bourne et al. (2007). Goodchild et al. (2012) estimated that the SICCT as interpreted in the RBCT, which includes the severe interpretation, had a specificity of 99.87%.

To quantify the potential impact of imperfect diagnostic test specificity on the classification of ‘unconfirmed’ herd breakdowns, the standard epidemiological framework for false positive estimation were applied. The expected number of false positive herds (FP) in a tested population is a function of the total number of uninfected herds (N_uninfected)_ and the false positive rate (FPR) of the diagnostic test: FP = N_uninfected_ x FPR

The false positive rate is derived directly from the diagnostic specificity (Sp = 99.87%), representing the probability that a truly uninfected herd yields a positive test result: FPR = 1 - Sp = 1 – 0.9987 = 0.0013 or 0.13%

Because a subset of herds (n_confirmed_ = 882) was conclusively confirmed as true positives (see confirmed breakdowns in supplementary material) via postmortem examination, the maximum possible pool of truly uninfected herds is bounded by subtracting these confirmed cases from the total population (N = 2469):

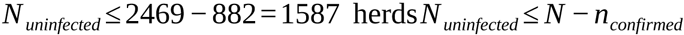

*FP* = 1587 *×* 0.0013 = 2 herds Substituting these parameters into the primary equation yields the total expected number of false positive herds across the entire trial:

Because any false positive herd must, by definition, fail to be confirmed upon postmortem examination, all expected false positives must reside within the unconfirmed cohort (*n*_unconfirmed_= 400). Consequently, out of the 400 positive skin test herds lacking postmortem confirmation (see unconfirmed breakdowns in supplementary data), only ≈ 2 are mathematically expected to be false positives. The remaining ≈ 398 unconfirmed herds represent true infections that were missed during postmortem inspection due to the known sensitivity limitations of gross postmortem pathology. Also the analysis assumes that all unconfirmed breakdowns were at the severe interpretation of the SICCT. The standard interpretation has a specificity of 99.98%, so this may overestimate the number of false positives.

### 2.3 Information Criteria and Model Fit Metrics

Model selection parsimony was evaluated using the corrected Akaike Information Criterion (AICc) alongside a custom formulation of the small sample size corrected Bayesian Information Criterion (BICc) (Ventura, Saulo & Leiva, 2019) for complex asymmetric contexts:

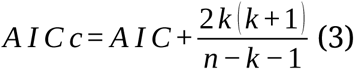

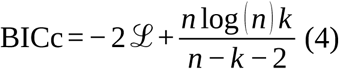

where ℒ is the maximized log-likelihood, *k* is the model’s degrees of freedom, and n represents the sample size based on the model residuals.

### 2.4 Permutation testing and null distribution architecture

To circumvent parametric assumptions and control for the highly structured nature of the experimental design, the significance of the treatment effect (*β*₁) was evaluated using non-parametric cluster-randomization permutation tests (Braun & Feng, 2001) over 10,000 successful iterations. Under the null hypothesis of no treatment effect, the treatment allocations within each matched triplet block are exchangeable. Following the established framework (Rosenbaum, 2020), the baseline environmental features, historical incidence rates, and asymmetric herd counts were kept fixed to preserve the true design matrix of the study. Randomly rearranging the treatment labels strictly within the matched triplet boundaries allowed construction of an empirical null distribution conditioned entirely on the observed structural features of the field sites.

Two alternative permutation frameworks were executed to diagnose the mathematical stability of the inference:

#### Model A (Unstudentized Coefficient Test)

The empirical null distribution was constructed using the raw regression coefficient 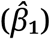 from each permuted iteration. The empirical p-value was computed as the proportion of shuffled iterations where the absolute null estimate exceeded or equalled the absolute observed estimate:

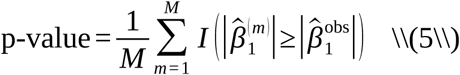

#### Model B (Studentized Wald test)

The empirical null distribution was constructed using the Wald Z-statistic 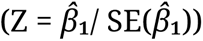 from each iteration. This framework standardizes the coefficient against shifts in standard error across random data partitions, providing an asymptotically pivotal test robust to localized variance inflation (Chung & Romano, 2013; Pauly, Brunner & Konietschke, 2015).

Crucially, in both frameworks, only the primary trial variable (treatment) was shuffled within its matched triplet boundary. Continuous baseline covariates remained fixed to their observed observations to preserve the structural covariance matrix of the data. Optimization health was rigorously filtered; any iteration failing to achieve clean mathematical convergence was discarded and logged.

### 2.5 Permutation leverage tracking and tail diagnostics

To mathematically diagnose which triplet blocks were driving the empirical null distributions, an automated leverage tracker within the 10,000-replicate permutation loop were implemented. For every shuffle where the simulated treatment estimate or Wald *Z*-score fell into the extreme empirical tail (*α ≤* 0.05), the exact composition of the randomized treatment assignments was decoded. An Over-Representation Ratio (ORR) was calculated for each cluster block by dividing its observed appearance frequency in the distribution’s tail by its mathematically expected uniform appearance under pure randomization. An ORR > 1.0 indicates that a specific triplet exerts disproportionate leverage over the test statistic’s variance boundaries. Further details are in the appendix.

### 2.6 Mathematical Formulation of Permutation Tail Leverage and the Over-Representation Ratio (ORR)

#### 2.6.1. Defining the Permutation Allocation Vector

Let the study design consist of *K* discrete, independent spatial blocks (triplets). Within each triplet *k* ∈ {1, 2, …, *K*}, let **x***_k_* define the localized vector of treatment assignments allocated to its constituent field sites. For a given permutation iteration *m* out of a total ***M*** successful shuffles (***M*** = 10,000), the global permutation state matrix ***X****_m_* is defined by the joint allocation:

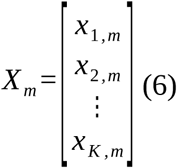

Because the random rearrangements are executed strictly within the matched triplet boundaries to preserve the true design matrix, the localized treatment vector **x***_k,m_* is drawn uniformly from the finite sample space of all structurally permissible local permutations Ω*_k_*. Under pure randomization, the probability of observing any specific local configuration *x_k_*\* remains uniform across iterations:

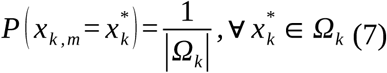

#### 2.6.2 Empirical Partitioning of the Statistical Tail

To isolate which local spatial blocks drive the boundaries of the test statistic’s variance, an objective conditional tail partition was executed on the empirical null distribution. Let *θ_m_* represent the simulated treatment estimate (Model A) or Wald Z-score (Model B) extracted from the *m*-th permutation replication. The extreme empirical tail space *Ψ_t_* is formally bounded using the *t*-th empirical quantile of the collected permutation distribution:

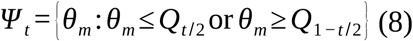

where *Q* satisfies the frequentist condition *P* (*θ* ∈ *Ψ_t_*) = *t*. For the diagnostic matrix executed in this study, the threshold is locked to the two-tailed 5th percentile extreme space (*t* = 0.05), yielding (0.025) area per tail). Let *M_Ψ_* = ∣*Ψ_t_*∣ represent the absolute number of permutation iterations falling within this partitioned tail space.

#### 2.6.3. Formulation of the Over-Representation Ratio (ORR)

Under the null hypothesis that the trial’s experimental units are structurally balanced and homogeneous, no single triplet’s localized assignment should disproportionately dictate whether the global test statistic lands in the tail space. Let I*_k_*_, *m*_ be a binary indicator variable showing whether the permuted treatment assignment for triplet *k* at iteration *m* matches its true, empirically observed field assignment *x_k_*_, obs_:

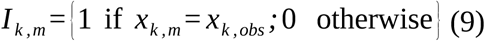

The empirical appearance frequency of the true configuration within the extreme tail space is defined as:

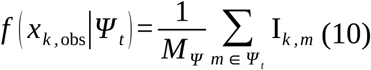

The global baseline expected frequency under pure, unconditioned uniform randomization across the entire permutation matrix is defined as:

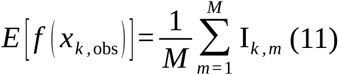

The Over-Representation Ratio (ORR*_k_*) for triplet *k* is formulated as the ratio of these two empirical profiles:

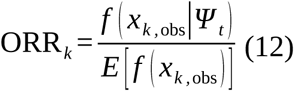

An ORR*_k_* of approximately 1.0 confirms that the localized treatment assignment within triplet *k* exerts no disproportionate leverage over the test statistic’s variance boundaries. Conversely, an ORR*_k_*> 1.0 demonstrates that a specific triplet’s observed configuration is structurally over-represented in the tail space.

### 2.7 Perturbation hypothesis model framework

To evaluate the boundary-expansion or “ perturbation” hypothesis, a parallel set of generalized linear mixed models evaluating disease incidence in herds flanking the culling areas (Neighbour_incidence) was specified. A random intercept for triplet was incorporated to account for unobserved regional baseline spatial heterogeneity. Exposure scaling properties was explicitly tested by contrasting a free log-linear reparametrisation, against a standard fixed exposure offset (offset (log (Neighbour_Herds *x* Timeatrisk))) across both nested null and treatment-bearing frameworks.

### 2.8. The binomial/logistic revision (Godfray et al. 2025)

Godfray et al (2025) stated “it would also be more appropriate to model the odds of a herd breakdown rather than the number that occur. Technically this implies fitting a binomial log-odds, rather than a Poisson log-linear, regression model, herd breakdown rather than the number that occur […]such an analysis, which has been independently refereed estimated the reduction of confirmed breakdowns as 17% (CI: 3% to 30%) and found that the reduction is significant at the standard 5% statistical level”.

The model proposed is a parametric logistic framework modelled via a binomial generalized linear model:

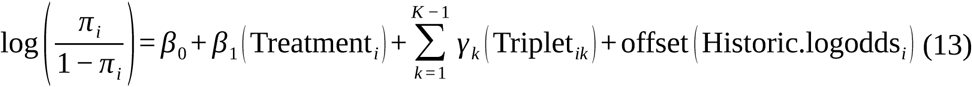

where the dependent variable is structured as a two-column matrix binding the number of successes (Incidence) and failures (Herds – Incidence). While this framework attempts to account for population variation, it enforces independent Bernoulli trial assumptions, implying that each herd represents a single independent outcome. Because the experimental time frame spans multiple years, some individual herds are subject to recurring, multiple breakdowns. Substantial numbers of the reported breakdowns in the RBCT are such recurrent breakdowns of the same herds. During the entire study period Donnelly et al (2007) reported 472 confirmed breakdowns among herds identified by VetNet as being inside the proactive trial (treatment) areas. These consisted of 245 herds that had a single confirmed breakdown, 74 herds had two confirmed breakdowns, and 25 herds that had more than two (3.16 on average). So approximately 29% of herds had multiple breakdowns, whilst about 50% of breakdowns were in herds suffering multiple breakdowns. The presence of large numbers of recurring counts completely invalidates the binomial distribution’s structural assumptions.

Nevertheless, the information-theoretic validity of this framework was evaluated in this paper by correcting Godfray’s model selection logic. While the 2025 review relied on unadjusted AICc profiles, a mathematically rigorous Quasi-AICc (QAICc) criteria using the variance inflation factor was implemented 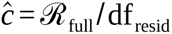 to evaluate model parsimony under observed overdispersion conditions:

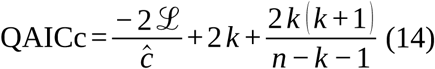

The binomial model was further analysed by introducing the time at risk, random effects and the complementary log-log link function.

## 3. Results

All statistical code, models and results are provided in the supplementary data file.

### 3.1 Parametric replication and diagnostic misspecification of the historical baseline

Replicating the historical fixed-effects Poisson formulation (model1) Donnelly et al. (2006) on the confirmed breakdown dataset reproduced a highly significant treatment effect (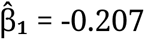, SE = 0.0729, Z = −2.835, p = 0.00458), seemingly providing strong evidence for a culling benefit.

Because the fixed-effects framework estimates a distinct intercept for every cluster block, it consumes 13 parameters on a dataset containing only 20 discrete observations, leaving just 7 degrees of freedom. When adjusting for this over parametrization via information criteria optimized for finite sample sizes, the model’s parsimony score drops away, causing the standard AIC of 142.29 to rise to a corrected AICc of 202.96. Furthermore, formal evaluation detected high residual dependency, throwing a critical autocorrelation warning (*p* < 0.001). Simulated quantile residual profiling via DHARMa exposed systemic violations of GLM assumptions, showing severe, significant quantile deviations and disperson (*p*_dispersion_ =0.0018, *p*_quantile_=0.011, Figure 1). This diagnostic layout evidences that the standard errors of the historical baseline are substantially deflated by unmeasured spatial autocorrelation, and the omission of the temporal exposure offset.

**Figure 1.**
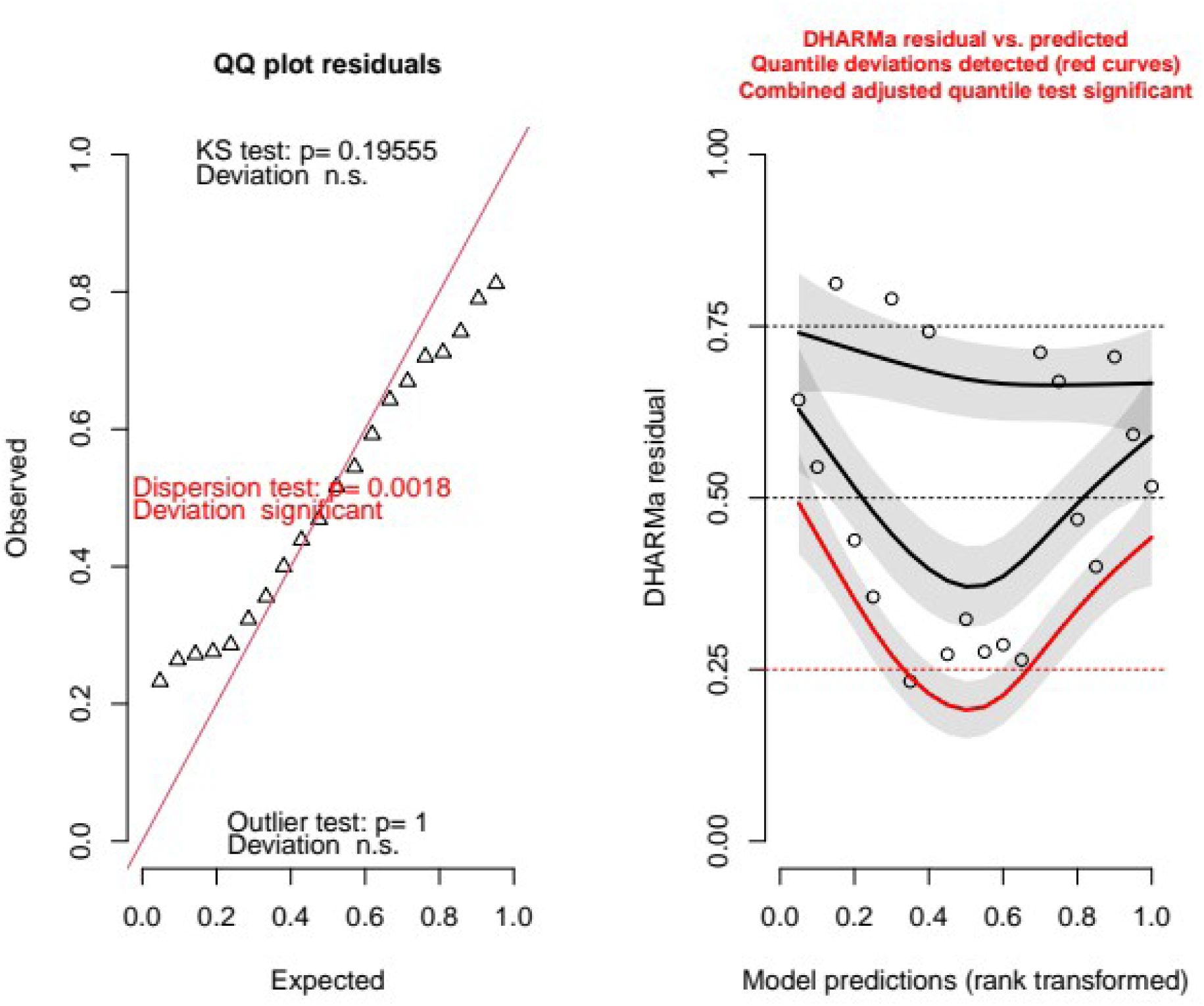
(Left) QQ plot of residuals of the original Donnelly et al (2006) Poisson model: the explicit dispersion test reveals substantial misspecification (p=0.0018), demonstrating a failure of the model’s parametric error structure to account for underlying data dispersion. (Right) Residuals vs. predicted values: Quantile deviations (solid black and red curves) are mapped against rank-transformed model predictions. The non-linear sinusoidal deviation from the nominal horizontal quantile lines (dashed red lines at 0.25, 0.50, and 0.75) indicates a significant structural misspecification of the model’s functional form. Combined with a parameter-to-data ratio of 13 parameters to 20 operational data points, these diagnostics mathematically demonstrate that the model overfits localized observation noise, creating the illusion of genuine epidemiological treatment pathways.

### 3.2. Generalized Mixed Model Specification

#### 3.2.1 “Confirmed breakdowns” analysis (inner area)

For the confirmed breakdown dataset, the observed treatment effect yielded an incidence rate ratio (IRR) of 0.846 (Z = -2.54) (model1r). Observed model selection criteria yielded highly conflicting signals. The full model and the null model (model2r) were essentially indistinguishable by AICc (ΔAICc = -0.05), presenting a statistical tie (Table 1). Conversely, BICc modestly favoured the null model (ΔBICc = +2.69).

**Table 1:** Treatment zone analysis results matrix.

| Dataset | Obs IRR | Obs Z | $\Delta\text{AICc}^*$ | $\Delta\text{BICc}^*$ | Model A $p$ | Model A 95% CI | Model B $p$ | Model B 95% CI |
| --- | --- | --- | --- | --- | --- | --- | --- | --- |
| Confirmed | 0.846 | -2.539 | -0.05 | +2.69 | 0.103 | [0.0945, 0.112] | 0.0624 | [0.0559, 0.0695] |
| Total | 0.925 | -1.052 | +3.11 | +5.85 | 0.459 | [0.445, 0.473] | 0.424 | [0.410, 0.438] |

\*In comparison to the null model

When subjected to non-parametric permutation, the seemingly strong observed Wald statistic was revealed to be a statistical anomaly. The diagnostic report across the 10,000 permutation replicates (coded as modelA or modelB) confirmed perfect mathematical stability, with zero convergence failures or hard code crashes recorded across all runs (N_valid_ = 10,000). Under modelA (raw coefficient), the treatment failed to achieve significance, yielding an empirical two-sided p-value of 0.103 (95% CI: 0.095, 0.112). When adjusting for standard error fluctuations via the studentized Wald approach (modelB), the empirical p-value shifted closer to the boundary but still markedly reduce the strength of evidence against Ho, yielding (p = 0.0625, CI: 0.0559, 0.0695).

#### 3.2.2 Total breakdown analysis (inner area)

When unconfirmed breakdowns were integrated into the GLMM (model1Tr)— incorporating all SICCT positive herds alongside their properly matched historical predictor (TotalHistoric)—any remaining indication of a treatment effect disappears. The observed IRR shifted toward neutrality at 0.925 (Z = -1.052) Both information criteria rejected the inclusion of the treatment vector, with ΔAICc = 3.11 and ΔBICc = 5.85 (Table 1) in favour of the nested null architecture (model2Tr). Consistent with this structural collapse, the non-parametric cluster-randomization permutations over 10,000 converged iterations yielded entirely flat empirical distributions, returning two-sided p-values for the treatment effect of 0.457 for Model A (95% CI: 0.445, 0.473) and 0.424 for Model B (95% CI: 0.410, 0.438). Correcting for the specificity of the SICCT (ie reducing the total breakdowns by 2 with 1000 random simulations across the data set) had a negligible effect on the analysis with the median effect size and *p* value being virtually identical (see supplementary data).

#### 3.2.3 Perturbation and random-effects scaling dynamics

For neighbouring areas, evaluating disease dynamics via a random-intercept framework inverted the exposure scaling verdict Mixed-effects models treating exposure as a free log-linear explanatory variable heavily outperformed standard fixed offset constraints. The preferred null explanatory configuration (model2P, AICc = 147.61) bettered the corresponding null fixed offset model (model4P, AICc = 153.56) by a substantial margin of ΔAICc = 5.95. This suggests that a transmission saturation effect operates outside the active culling perimeter identically to the cull area dynamics.

When evaluating the perturbation hypothesis within this preferred random-effects framework, the information criteria favoured the null model (modelP2). Incorporating the primary treatment variable (modelP1) resulted in an information tie under AICc (AICc = 147.85, ΔAICc = 0.23). Conversely, the BICc penalized the addition of the treatment vector, yielding a ΔBICc = 2.97 in favour of the nested null architecture (BICc_Full_ = 159.34, BICc_Null_ = 156.37) (Table 2).

**Table 2:** Neighbouring area analysis results matrix.

| Dataset | Obs IRR | Obs Z | $\Delta\text{AICc}^*$ | $\Delta\text{BICc}^*$ | Model A $p$ | Model A 95% CI | Model B $p$ | Model B 95% CI |
| --- | --- | --- | --- | --- | --- | --- | --- | --- |
| Confirmed_Neighbour | 1.219 | 2.149 | +0.23 | +2.97 | 0.0640 | [0.0574, 0.0711] | 0.0756 | [0.0684, 0.0833] |
| Total_Neighbour | 1.116 | 1.340 | +2.46 | +5.20 | 0.246 | [0.235, 0.259] | 0.245 | [0.233, 0.257] |

\*In comparison to the null model

For the confirmed neighbouring area breakdown dataset, the observed treatment variable returned an apparent risk inflation (IRR = 1.219, Z = 2.149) However, this parametric signal appears to be a cluster-dependent artifact under exact permutation. With Model A (raw coefficient), the perimeter effect failed to achieve statistical significance (*p* = 0.0640, 95% CI: [ 0.0574, 0.0711 ]), a finding echoed by the studentized Wald test under Model B (*p* = 0.0756, 95% CI: [ 0.0684, 0.0833 ]) (Table 2).

When diagnostic miss classifications were corrected by analysing the total neighbouring area dataset (modelPT1 and model PT2), including unconfirmed cases in the prior three year period, the statistical signal disappeared as background noise. The observed perimeter IRR shifted toward neutrality (IRR = 1.116, Z = 1.340) Both information criteria favoured the nested null architecture (ΔAICc = +2.46, ΔBICc = +5.20). Mirroring this informational rejection, the empirical null distributions flattened completely, yielding highly non-significant empirical *p*-values of 0.246 for Model A (95% CI: [0.235, 0.259]) and 0.245 for Model B (95% CI: [0.233, 0.257])

### 3.3 Structural unbalance and cluster leverage diagnostics

The permutation leverage tracking verified that the randomized design of the trial was unbalanced, with the empirical null distribution resting mainly on a structural anomaly within isolated clusters. For example, out of the 10 randomized blocks, triplet C (ORR=1.59) and triplet A (ORR=1.44) exhibited substantial over-representation in the significance tails of the shuffled runs (Table 3; Figure 2).

**Figure 2:**
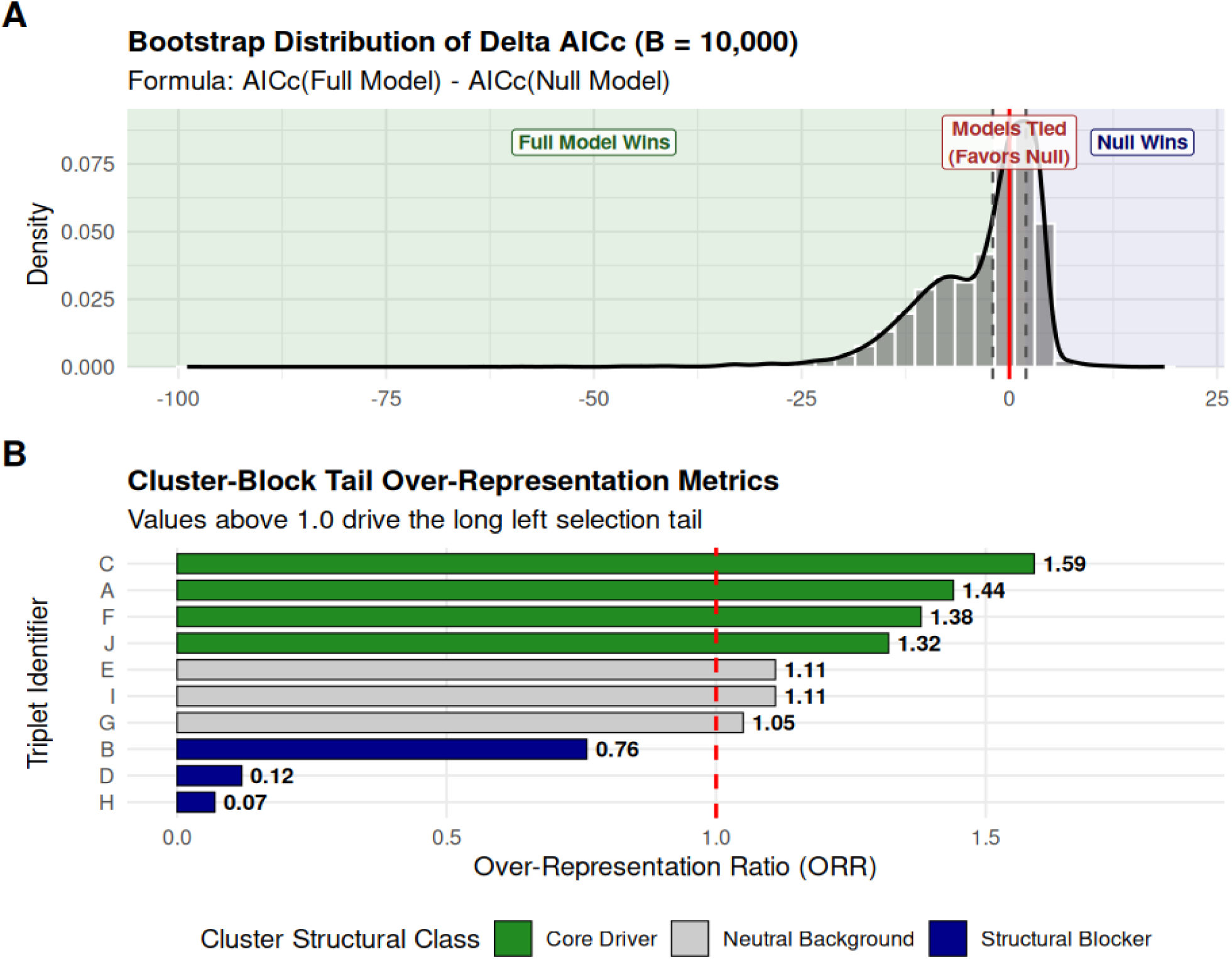
bootstrap distribution of DAICc of the full model (model1r) – the null model (model2r)

**Table 3:** Cluster leverage diagnostics. Table Note: Summary of cluster block leverage metrics within the 10,000-replicate empirical 5% left tail. Values highlighted for triplets C and A demonstrate structural unbalance, where random shuffles of these two clusters dictate the significance boundaries of the entire trial.

| Triplet ID | Avg Appearance In Tail | Expected Appearance | Over-Representation Ratio |
| --- | --- | --- | --- |
| C | 1.62 | 1.02 | 1.59 |
| A | 1.44 | 1.00 | 1.44 |
| F | 1.40 | 1.01 | 1.38 |
| J | 1.31 | 0.99 | 1.32 |
| E | 1.11 | 1.00 | 1.11 |
| I | 1.12 | 1.01 | 1.11 |
| G | 1.05 | 1.00 | 1.05 |
| B | 0.75 | 0.99 | 0.76 |
| D | 0.12 | 0.99 | 0.12 |
| H | 0.07 | 0.99 | 0.07 |

Conversely, non-responsive clusters like triplet H (ORR= 0.07) and triplet D (ORR = 0.12) were virtually absent from the tail thresholds. These metrics reveal that the global test statistic is highly sensitive to the specific treatment labels assigned within these two blocks, providing a clear mathematical foundation for identifying structural unbalance in the field design without violating the strict fixed constraints of the study matrix. This diagnostic grid strongly suggests that when the block effects are omitted (model1i), the apparent treatment effect disappears because the severe, non-uniform variance inflation anchored within triplets A and C is diluted across the background noise of the balanced, non-responsive trial design.

### 3.4 Information-theoretic reversal of the binomial framework

Replicating the binomial specifications proposed by Godfray et al (2025) highlights the fragility of relying on unadjusted model selection tools. Running standard, AICc criteria replicates Godfray’s original hierarchy, where the treatment model containing a fixed historical offset (modelB3, AICc = 173.58) outperforms the nested Null model (modelB4, AICc = 173.87) by a negligible D of -0.29. Based on this profile, fitting a quasibinomial adjustment (modelB3q) yields a marginally significant treatment effect (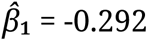, SE = 0.124, *t* = -2.363, *p* = 0.0424).

However, the summary profile reveals that this significance rests on a corrupted variance matrix, with the empirical dispersion parameter inflating to a 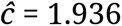. When the correct information criteria are applied by propagating this variance inflation factor through a proper QAICc parameterization, the model selection direction reverses. The treatment model’s score drops to QAICc = 132.81, while the nested Null model (ModelB4) achieves a far improved score of 125.11. This generates a definitive information difference of ΔQAICc = 7.70 in favour of the Null model. Under information-theoretic standards, a positive D of this magnitude demonstrates that incorporating the treatment parameter provides no real explanatory utility. Further analysis incorporating time at risk and random effects models is reported in the supplementary data file, but do not change the conclusion.

### 3.5 Sensitivity to Cluster Intercept Omission

To isolate the structural mechanism behind the culling metrics, an isolated parameter check was executed by completely removing the spatial blocking variable triplet from the generalized Poisson regression (model1i). This structural omission caused the treatment parameter to become nominally non-significance (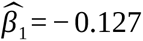, SE = 0.088, *Z* = *−* 1.441, *p* = 0.150), while the residual matrix achieved formal independence (p = 0.440). Crucially, even when the model was stripped of all spatial cluster intercepts, the empirical scaling parameter for the exposure variable remained at 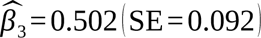.

### 3.6 Deconstructing the saturation artifact: recurrence vs density dependence

The discovery that the empirical exposure parameter stabilizes at 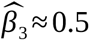 across both the preferred mixed models and the fixed effect model (Model1i) allows potential resolution of the underlying mechanism of the apparent transmission saturation effect. Historically, information-theoretic support for a free explanatory exposure term over a rigid fixed offset may be hypothesized to reflect ecological constraints. While ecological factors may coexist, the primary driving force behind this sub-linear scaling is operational and statistical: it is simply a possible consequence of herd-level disease recurrence.

Because the trial data tracks herds over an extended multi-year timeline, larger herds with higher baseline risks are susceptible to recurring breakdown events. The event rate undergoes log-linear compression as the outbreak is managed and cleared. By freeing the exposure parameter, the generalized Poisson model accommodates this recurrence by fitting a compressive slope of ≈ 0.5.

This mechanical reality invalidates both the Donnelly fixed effects Poisson model and the Godfray binomial revision. The Donnelly fixed effect model has 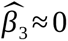 which would suggest that breakdowns varied little with herd numbers within a triplet area and is additional evidence for misspecification. On the other hand this can not be rectified by assigning a rigid offset (*β* = 1, as it would artificially inflate the expected background risk in high-exposure cohorts, forcing the excess count variance into the residual matrix and triggering the profound autocorrelation (p < 0.001) and quantile deviations detected in DARMa profiling. Similarly, the Godfray binomial framework is biologically and mathematically incorrect; it treats the herds as a static vector of independent Bernoulli trials, a structural assumption that contrasts with the reality of multiple, recurring breakdowns within a single herd unit. Despite this further binomial models were considered which included block random effects and a complementary log-log link to enable modelling time at risk. These are detailed in the supplementary data file, and none were able to legitimise a treatment effect.

When these historical misspecifications are removed, the full inferential architecture of the trial unifies. The treatment effect is absent. The apparent cull area benefits and neighbouring area “negative effects” reported across two decades of literature are revealed to be a interconnected chain of mathematical artifacts: caused by a recurrent count system, over-filtering data based on imperfect post-mortem sensitivity, and relying on parametric tests vulnerable to high-leverage cluster anomalies.

## 4 Discussion

A striking finding of this re-examination is the sub-linear exposure slope (*b* ≈ 0.50) estimated across all free-covariate models, a signature that remained unchanged when evaluating both confirmed breakdowns and total incidence. Biologically, infectious disease risk is expected to scale linearly (*b*= 1.0) with population density and exposure time. The model degradation (overdispersion and quantile residual deviations) that occurs when enforcing a strict offset reveals a fundamental mismatch between standard count-model assumptions and the true biological dynamics of the trial.

This mismatch may be driven by herd-level recurrence inside the trial areas. The limited data of herd-level recurrence documented (Donnelly et al., 2007; Karolemeas et al., 2012) did not report the data by triplet so it was not possible to incorporate this data into the present analysis. However, it is clear that a subset of chronic herds recurrently broke down regardless of marginal changes in herd size or overall time-at-risk. By pooling these recurrent, highly correlated within-herd events into a simple global count, historical modelling frameworks conflated widespread transmission with localized chronic recurrence. This distorted the exposure parameters and left the model incapable of adequately separating true intervention effects from background recurrence noise.

The initial parametric significance observed in the confirmed breakdowns dataset (Z = -2.539) represents an inferential anomaly driven by extreme data leverage within localized cluster blocks rather than a systemic, population-wide treatment effect. Historical analytical frameworks were trapped in a structural Catch-22: retaining the trial’s matched triplets as fixed dummy variables consumed excessive degrees of freedom from a sparse 20-row dataset, resulting in severe over fitting and profound residual autocorrelation. Conversely, completely omitting the spatial blocking variable dilutes the localized, high-variance signals of outlying clusters (specifically triplets A and C) across the background noise of non-responsive blocks, causing the treatment parameter to collapse (*p* = 0.150). By applying a random-intercept mixed-effects specification, our architecture resolves this statistical paradox. This approach estimates a single variance component 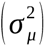 to protect critical degrees of freedom while strictly respecting the restricted randomization block design. That the treatment effect fails to achieve significance under this parsimonious random-intercept model (*p*_Model B_ = 0.0624), the omitted-block model (*p* = 0.150), and exact non-parametric cluster-constrained permutations (*_p_*_Model A_ = 0.103) consistently demonstrate that the apparent efficacy of proactive culling is an artifact of regional imbalance, incapable of withstanding robust structural verification.

These findings provide unified evidence that badger culling lacks robust verification required to guide national disease control policy. This is in strict contrast to the assertion made by Donnelly et al (2006) who stated: “This result was consistent across ten proactive culling areas, each paired with a survey-only area”.

The structural collapse of the historical baselines under modern diagnostic profiling provides the mathematical explanation as to why historical analyses concluded large intervention effects. The model implemented by Donnelly et al (2006) (model1) suffers from over parametrization, consuming 13 parameters on a dataset containing only 20 discrete observations. This results in the standard AICc to penalize it heavily (AICc = 202.96). By forcing a rigid Poisson error distribution on data featuring strong spatial and temporal dependencies, the model traps local regional clustering within its residual matrix, triggering the profound autocorrelation, quantile deviation and dispersion detected in the diagnostics.

The failure of the Godfray et al. (2025) Strategy Review model under modern statistical profiling reveals a deeper structural issue that shapes the entire RBCT evaluation. The review’s insistence on a significant culling effect (*p* = 0.042) stems from a compounding mathematical contradiction: using a standard binomial distribution for model selection while switching to a quasibinomial model for inference. By executing model selection on a standard binomial framework that ignores overdispersion, the original analysis selected a parameter-heavy treatment model (modelB3). Once overdispersion is correctly integrated into the selection step using the empirical variance inflation factor (ĉ = 1.94), QAICc heavily rejects the treatment model, favouring the nested null model by a large margin of ΔQAICc = 7.70.

Beyond this mathematical contradiction, the binomial specification suffers from an unresolvable biological flaw. Defining the trial outcomes as a series of binomial successes and failures assumes that each herd is a single binary unit with a single episode; breaking down only once. In a long-term field study, herds can experience multiple, recurring breakdown events, some of which may be due to recrudescence rather than fresh infection. This recurrent counting structure invalidates the independent Bernoulli trial assumption that defines the binomial distribution. Forcing data with recurring counts into a binomial matrix distorts the model’s denominator, resulting in severe overdispersion that standard quasi-likelihood adjustments can only patch over, not fix. Likewise other models with the binomial architecture can not repair this issue.

The definitive evidence that the marginal parametric culling effect is an artifact of human detection bias emerges when these models are reconciled with the diagnostic parameters of the bovine tuberculosis tracking pipeline. Because the primary live-animal diagnostic skin test (SICCT) operates with an exceptional field specificity approaching 100%, the unconfirmed herd breakdowns cannot be dismissed as false positives; they represent true biological infections. The systemic divergence between live-animal reactor status and post-mortem confirmation is driven entirely by the severe sensitivity limitations of standard slaughterhouse macro-necropsy (46%–79%), rendering the administrative line separating ‘confirmed’ from ‘unconfirmed’ statuses highly volatile and dependent on operational noise, such as carcass throughput speeds and individual pathologist visual acuity. Biologically, herds with unconfirmed breakdowns represent the front line of active transmission and early-stage infections before advanced gross pathology develops. Critically, there is no plausible epidemiological mechanism by which wildlife host removal could selectively suppress only those infections that progress to macroscopic lesions while leaving latent infections unaffected. The corrected side-by-side analysis demonstrates the exact opposite: when diagnostic misclassification bias is eliminated by restoring these true biological infections to the total reactor dataset, any trace of a culling effect disappears across all statistical frameworks (p_Model A = 0.459, ΔBICc = +5.85), and information criteria overwhelmingly favour nested null architectures. The historical claim of culling efficacy is therefore revealed to be an illusion of abattoir sensitivity tracking observation noise rather than a genuine intervention response.

The side-by-side analysis of the breakdown incidence in neighbouring areas provides the final critical link in RBCT evaluation, completing a symmetric statistical story. Historically, wildlife management policies have been contested based on the perturbation effect hypothesis and the conclusions of the RBCT experiment (Bourne et al., 2007). The non-parametric cluster-permutations indicate no ‘positive’ or ‘negative’ treatment effects are measurable.

Inside the culling boundary, any apparent positive treatment effect fails exact permutation (*_p_*_Model A_ = 0.103) and completely disappears when unconfirmed breakdowns are included (*_p_*_Model A_ = 0.459), ΔBICc = 5.85). Outside the culling boundary, the same statistical issue occurs. The apparent neighbouring area risk inflation observed in confirmed cases lacks systemic stability and falls short of non-parametric significance under both unstudentized (*p* = 0.0640) and studentized (*p* = 0.0756) logic. When human observation limits and slaughterhouse necropsy sensitivity biases (46%--79%) are bypassed by restoring ‘unconfirmed’ SICCT reactors to the total neighbouring areas dataset, the perturbation effect vanishes (*_p_*_Model A_ = 0.246, *_p_*_Model B_ = 0.245), while BICc criterion favours the null model (ΔBICc = 5.20).

This mathematical harmony between the cull and neighbouring areas reveals that badger culling did not reduce herd incidents of bovine tuberculosis—rather it is statistical noise. When evaluated using non-parametric resampling tools that protect against high-variance cluster leverage, the badger intervention is revealed to be epidemiologically inert across the RBCT experiment.

Vial, Johnston & Donnelly (2011) applied a Cox proportional hazards model to RBCT data, which revealed that the effect of proactive culling on the time to first confirmed bTB breakdown was not statistically significant (*p* = 0.08). However the Poisson regression model (model 1) was favoured over the Cox model for inference in respect of any treatment effect. Nevertheless, this finding indicates that isolating independent initial events, rather than pooling recurrent herd breakdowns, results in a loss of treatment effect significance. This previous finding is consistent with and reinforces the conclusions of the present study.

Mills, Woodroffe & Donnelly (2024a, 2024b) concluded that the Poisson regression model (model 1) was the most robust approach, but failed to uncover the issues documented in this analysis. Torgerson et al. (2024) suggested, by parsimony of competing models, that treatment was unlikely to have an effect on incidence of bTB in cattle. However the most parsimonious model removed triplet from the analysis, which undermined the block design of the trial. The present study confirms this previous finding whilst retaining the block design. In the Godfray et al. (2025) review it was concluded that an offset was essential to control for the numbers of herds in each block. To deal with this issue, they proposed the logistic regression model which has been shown to be flawed in the present study. Bayesian model architecture with minimally informative priors did not alter the conclusions of the relevant frequentist approaches (Mills, Woodroffe & Donnelly, 2024a, 2024b; Torgerson et al., 2024, 2025). The present study is the only study to introduce a random effects to maintain the block design, hence increasing the numbers of degrees of freedom.

When statistical models directly inform massive public expenditure and agricultural interventions, relying on a binary, uncorrected *p* < 0.05 threshold introduces immense financial and operational liability. In this study, the conflict between localized parameter stability tests and global model-comparison bootstraps, combined with the definitive null results of the total incidence analysis, represents the classic signature of an underpowered data structure. The data contains localized signals in restrictive subsets, but completely lacks the statistical volume and consistency required to generate actionable evidence for large-scale deployment.

Consequently, it is concluded there is insufficient evidence to support the hypothesis that proactive badger culling can reduce the incidence of confirmed herd breakdowns of tuberculosis in cattle. The incorporation of the full diagnostic disease burden — including unconfirmed skin-test reactors — fundamentally shifts the baseline narrative of this study. The resulting data structure does not merely present a case of underpowered ‘insufficient evidence’ for an intervention effect. Rather, the model metrics provide strong, affirmative evidence that any true proactive culling treatment effect is highly unlikely. This is demonstrated by two distinct mathematical behaviours. First, the introduction of the treatment variable to the corrected dataset of all breakdowns explicitly degrades global model fit, resulting in DAICc penalty of 3.11 and allocating a 82.5% of the Akaike weight directly to the null framework. Second, the estimated intervention effect size attenuates to a negligible (model1Tr, b = -0.077, p = 0.293) that sits entirely within the realm of random baseline noise. Consequently, when diagnostic under-reporting is corrected, the data actively supports the null hypothesis, demonstrating that an effective population-level intervention response is highly improbable within this trial framework.

### Policy implications

Advancing to a full-scale policy roll out under an empirical risk threshold that favours the null model carries an unacceptably high risk of spending vast public resources for zero net epidemiological benefit. The analysis reported here suggests that the perturbation effect is also a statistical artifact.

The most responsible policy translation of these findings is to conclude that badger culling should play no role in the control of tuberculosis in cattle. This recommendation is the same conclusion of the authors of the RBCT (Bourne et al., 2007). However this earlier recommendation was due to the possibility of the perturbation effect abrogating the culling effect. This recommendation was overruled by the King report (King, 2007) which concluded any perturbation effect could be avoided by culling very large numbers of badgers over wider areas: “In our view a programme for the removal of badgers could make a significant contribution to the control of cattle TB in those areas of England where there is a high and persistent incidence of TB in cattle, provided removal takes places alongside an effective programme of cattle controls”.

A widespread badger culling programme commenced in 2013. Analyses of this culling programme (Langton, Jones & McGill, 2022) strongly suggested that mass proactive badger culling has had no detectable effect on the incidence of tuberculosis in cattle herds.

Godfray et al. (2025) in the latest evidence review stated “our conclusion is that the analysis of confirmed incidents in the RBCT provides weaker evidence for a positive effect of culling than its first analyses suggested […]. Government policy is to end culling but even were it to continue the RBCT now provides limited (if any) insights into the design and likely value of including culling in a control programme “.

Despite this, the RBCT continues to be a key reference cited in the latest DEFRA policy documents (Cross et al. 2026) as evidence that badger interventions are a legitimate strategy for the control of TB in cattle.

Cox and Donnelly, using the RBCT as an example, argued that “requiring independent replication of specific statistical analyses as a general check before publication seems not merely unnecessary but a misuse of relatively scarce expertise” and “the purpose of providing the full data was not primarily to encourage repetition of our own analyses, [but] to encourage […] the formulation and investigation of new questions” (Cox & Donnelly, 2010). However, the presence of highly significant residual autocorrelation and severe quantile dispersion anomalies across the fixed-effects formulation in the Poisson model proposed by Donnelly et al. (2006) underscores a critical vulnerability regarding the structural validity of historical RBCT analytical frameworks. True reproducibility requires not only the rerun of computational workflows but the strict satisfaction of the mathematical assumptions underpinning the chosen statistical estimators. When a traditional parametric model is applied to spatially and temporally clustered data without accounting for structural dependencies, the assumption of independent and identically distributed errors is explicitly violated. This over fitted architecture inherently mistakes observation noise for real epidemiological trends. Data-dependent analysis and structural misspecification remain primary drivers of the wider replication crisis (Spanos, 2024), proving that routine independent technical audits are a mandatory safeguard for public policy.

In epidemiological modelling, ignoring this violation induces artificial deflation of standard errors and a corresponding overstatement of statistical precision (type I error inflation). Consequently, historical beliefs regarding the efficacy of badger culling interventions rest on unstable inferential foundations. Because model selection metrics like AICc assume that the underlying probability distribution is correctly specified, an unmodeled correlation structure can distort model weights and lead to the premature retention or exclusion of critical risk factors.

Reconsidering this analysis through a generalized linear mixed model framework is a prerequisite for robust reproducibility. Likewise bootstrap analysis supplements selection metrics to identify the most robust model.

Some may prefer to identify the RBCT now as an inconclusive experiment, due to the limitations of its design, many of which have become apparent in hindsight. However, failure to correct the diagnostic shortcomings of the RBCT analysis over the last two decades has propagated long-term systemic inferential biases into public health and wildlife management policies. Consequently, the culling of over 250,000 largely healthy badgers since 2013 in England has had no demonstrable effect on in the incidence of bTB in cattle herds. It can be considered a misdirected use of resources, and, like mass badger vaccination, a speculative and extreme intervention with no credible scientific foundation, economic, public health or animal welfare justification.

## Data Availability Statement

All relevant raw epidemiological data, including herd incidence counts, total skin-test incidence counts, treatment allocations, historical baseline indices, and exposure vectors (herd sizes and time-at-risk) across both the proactive culling zones and neighboring spillover zones, are included in full within the manuscript submission (specifically as **Supplementary Data File 1**).

## References

Bourne, F.J., Donnelly, C.A., Cox, D.R., Gettinby, G., McInerney, J.P., Morrison, W.I. (2007) Bovine TB: The scientific evidence: A science base for a sustainable policy to control TB in cattle. An epidemiological investigation into bovine tuberculosis: Final Report of the Independent Scientific Group on Cattle TB. Department for Environment, Food and Rural Affairs. London.

Brooks, M.E., Kristensen, K., Van Benthem, K.J., Magnusson, A., Berg, C.W., Nielsen, A., Skaug, H.J., Mächler, M. & Bolker, B.M. (2017) glmmTMB balances speed and flexibility among packages for zero-inflated generalized linear mixed modeling. The R Journal, 9(2), 378–400.

Braun, T.M. & Feng, Z. (2001) Optimal permutation tests for the analysis of group randomized trials. Journal of the American Statistical Association, 96, 1424– 1432.

Chung, E. & Romano, J.P. (2013) Exact and asymptotically robust permutation tests. The Annals of Statistics, 41, 484–507.

Cox, D.R. & Donnelly, C.A. (2010) Commentary. Biostatistics, 11, 381–382.

Donnelly, C.A., Woodroffe, R., Cox, D.R., Bourne, F.J., Cheeseman, C.L., Clifton-Hadley, R.S., Ghani, A.C., Gettinby, G., Gilks, P., Jenkins, H., Johnston, W.T., Le Fevre, A.M., McInerney, J.P. & Morrison, W.I. (2006) Positive and negative effects of widespread badger culling on tuberculosis in cattle. Nature, 439, 843–846.

Donnelly, C.A., Wei, G., Johnston, W.T., Cox, D.R., Woodroffe, R., Bourne, F.J., Cheeseman, C.L., Clifton-Hadley, R.S., Gettinby, G., Gilks, P., Jenkins, H., Le Fevre, A.M., McInerney, J.P. & Morrison, W.I. (2007) Impacts of widespread badger culling on cattle tuberculosis: concluding analyses from a large-scale field trial. International Journal of Infectious Diseases, 11, 300–308.

Godfray, H.C.J., Donnelly, C.A., Hewinson, R.G., Winter, M. & Wood, J.L.N. (2018) Bovine TB strategy review. Department for Environment, Food and Rural Affairs (DEFRA), London, UK.

Godfray, H.C.J., Hewinson, R.G., Silverman, B., Winter, M. & Wood, J.L.N. (2025) Bovine TB strategy review update 2025. Department for Environment, Food and Rural Affairs (DEFRA), London, UK.

Goodchild, A.V., Downs, S.H., Upton, P., Wood, J.L.N. & de la Rua-Domenech, R. (2015) Specificity of the comparative skin test for bovine tuberculosis in Great Britain. Veterinary Record, 177, 258–258.

Hartig, F. (2024) DHARMa: Residual Diagnostics for Hierarchical (Multi-Level / Mixed) Regression Models. R package version 0.4.7.

Karolemeas, K., Donnelly, C.A., Conlan, A.J.K., Mitchell, A.P., Clifton-Hadley, R.S., Upton, P., Wood, J.L.N. & Andrew, A.J. (2012) The effect of badger culling on breakdown prolongation and recurrence of bovine tuberculosis in cattle herds in Great Britain. PLOS ONE, 7, e51342.

King, D. (2007) Bovine Tuberculosis in cattle and badgers. Department for Environment, Food and Rural Affairs (DEFRA), London, UK.

Langton, T.E.S., Jones, M.W. & McGill, I. (2022) Analysis of the impact of badger culling on bovine tuberculosis in cattle in the high-risk area of England, 2009– 2020. Veterinary Record, 190, e1384.

Mills, C.L., Woodroffe, R. & Donnelly, C.A. (2024a) An extensive re-evaluation of evidence and analyses of the Randomised Badger Culling Trial (RBCT) I: Within proactive culling areas. Royal Society Open Science, 11, 240385.

Mills, C.L., Woodroffe, R. & Donnelly, C.A. (2024b) An extensive re-evaluation of evidence and analyses of the Randomised Badger Culling Trial II: In neighbouring areas. Royal Society Open Science, 11, 240386.

Nuñez-Garcia, J., Downs, S.H., Parry, J.E., Abernethy, D.A., Broughan, J.M., Cameron, A.R., Cook, A.J., de la Rua-Domenech, R., Goodchild, A.V., Gunn, J., Hanic, M., Hayton, A., Holliman, A., More, S.J., Nicholas, R.A., Palmer, S., Sharpe, M., Smith, G.C., Upton, P., Vordermeier, H.M. & Watson, E.N. (2018) Meta-analyses of the sensitivity and specificity of ante-mortem and post-mortem diagnostic tests for bovine tuberculosis in the UK and Ireland. Preventive Veterinary Medicine, 153, 94–107.

Pascual-Linaza, A.V., Gordon, A.W., Stringer, L.A. & Menzies, F.D. (2017) Efficiency of slaughterhouse surveillance for the detection of bovine tuberculosis in cattle in Northern Ireland. Epidemiology and Infection, 145, 995–1005.

Pauly, M., Brunner, E. & Konietschke, F. (2015) Asymptotic permutation tests in general factorial designs. Journal of the Royal Statistical Society: Series B (Statistical Methodology), 77, 461–473.

R Core Team (2026) R: A language and environment for statistical computing. R Foundation for Statistical Computing, Vienna, Austria.

Rosenbaum, P.R. (2020) Design of Observational Studies, 2nd edn. Springer, Cham, Switzerland.

Spanos, A. (2024) Revisiting the replication crisis and the untrustworthiness of published empirical evidence: a severe testing perspective. Journal of Risk and Financial Management, 17(1), 35.

Torgerson, P.R., Hartnack, S., Rasmussen, P., Lewis, F. & Langton, T.E.S. (2024) Absence of effects of widespread badger culling on tuberculosis in cattle. Scientific Reports, 14, 16326.

Torgerson, P.R., Hartnack, S., Rasmussen, P., Lewis, F.I., O’Donnell, P. & Langton, T.E.S. (2025) Randomised Badger Culling Trial—no effects of widespread badger culling on tuberculosis in cattle: comment on Mills, Woodroffe and Donnelly (2024a, 2024b). Royal Society Open Science, 12, 241609.

Ventura, M., Saulo, H., Leiva, V. & Monsueto, S. (2019) Log-symmetric regression models: information criteria and application to movie business and industry data with economic implications. Applied Stochastic Models in Business and Industry, 35, 963–977.

Vial, F., Johnston, W.T. & Donnelly, C.A. (2011) Local cattle and badger populations affect the risk of confirmed tuberculosis in British cattle herds. PLOS ONE, 6, e18058.

