## Supplementary file 2 for "A Computational Re-evaluation of Spatial Trials for Zoonotic Tuberculosis Control: Model Misspecification, Diagnostic Misclassification, and the Illusion of Wildlife Culling Efficacy": Suplementary_File_2.R

```
# ==============================================================================
#
# Randomized badger cull trial
# ==============================================================================

library(readxl)
RBCT <- read_excel("RBCT.xlsx")
library(MASS)
library (MuMIn)
library(AICcmodavg)
library(glmmTMB)
library(DHARMa)
library(performance)

RBCT$Triplet<-as.factor(RBCT$Triplet)
RBCT$Triplet = relevel(RBCT$Triplet, ref="J")


#BICc (Ventura et al 2019)
#Ventura, M., Saulo, H., Leiva, V., Monsueto, S., 2019. Log-symmetric regression models: information 
#criteria and application to movie business and industry data with economic implications.
#Applied Stochastic Models in Business and Industry 35, 963–977

BICc<-function (model) 
{
  LL <- logLik(object = model)
  df <- attr(LL, "df")
  n <- length(residuals(model))
  c(-2 * LL + n*log(n)*df/(n-df-2))
}

#model 1 is the original poisson regression from Donnelly et al, RBCT and comparisons when treatment
#is removed and/or exposure is coded as an offset.
model1<-glm(Incidence~Treatment+Triplet+log(Historic)+log(Herds),poisson, data=RBCT)
model2<-glm(Incidence~Triplet+log(Historic)+log(Herds),poisson, data=RBCT)
model3<-glm(Incidence~Treatment+Triplet+log(Historic)+offset(log(Herds)),poisson, data=RBCT)
model4<-glm(Incidence~Triplet+log(Historic)+offset(log(Herds)),poisson, data=RBCT)

#Model4 has the lowest AICc
unlist(lapply(list(model1, model2, model3, model4), AICc))
unlist(lapply(list(model1, model2, model3, model4), BICc))
aictab(list(model1,model2,model3,model4))

#All 4 models have issues with autocorrelation of residuals
check_autocorrelation(model1)
check_autocorrelation(model2)
check_autocorrelation(model3)
check_autocorrelation(model4)

#Model 1, the preferred model in Donnelly et al. 2006 has issues with residuals
sim1<-simulateResiduals(model1, n=10000)
par(mfrow = c(1, 2), cex = 0.8, mar = c(5, 4.5, 4, 3))
plotQQunif(sim1)
plotResiduals(sim1)
par(mfrow = c(1, 1), cex = 1.0, mar = c(5, 4, 4, 2))
testQuantiles(sim1)
#Random effects models maintain block effects (Triplet) and increase degrees of freedom.
#Models are coded as a generalized poisson to control for over or under dispersion 
#Each triplet has a unique time at risk, so time at risk is now included in the exposure variable
#Exposure is number of herds x time at risk during the experimental period
model1r<-glmmTMB(Incidence~Treatment+(1|Triplet)+log(Historic)+log(Herds*Timeatrisk),genpois, data=RBCT)
model2r<-glmmTMB(Incidence~(1|Triplet)+log(Historic)+log(Herds*Timeatrisk),genpois, data=RBCT)
model3r<-glmmTMB(Incidence~Treatment+(1|Triplet)+log(Historic)+offset(log(Herds*Timeatrisk)),genpois, data=RBCT)
model4r<-glmmTMB(Incidence~(1|Triplet)+log(Historic)+offset(log(Herds*Timeatrisk)),genpois, data=RBCT)

#Model1r and Model2r are virtually indistinguishable in AICc values.
#Model 2r has the lowest BICc
unlist(lapply(list(model1r, model2r, model3r, model4r), AICc))
unlist(lapply(list(model1r, model2r, model3r, model4r), BICc))
aictab(list(model1r,model2r,model3r,model4r))

#Model3r have issues with autocorrelation of residuals
check_autocorrelation(model1r)
check_autocorrelation(model2r)
check_autocorrelation(model3r)
check_autocorrelation(model4r)

#Model1r has a treatment effect
summary(model1r)

#Model1r and model2r are further analysed by permutation tests
#See later

#Model1i, remove triplet from the analysis, treatment effect disappears
#But loses block design
#Note Beta for exposure = 0.50
model1i<-glmmTMB(Incidence~Treatment+log(Historic)+log(Herds*Timeatrisk),genpois, data=RBCT)
summary(model1i)

#No issues with autocorrelation of residuals, but quantile deviations detected
check_autocorrelation(model1i)
plot(simulateResiduals(model1i))

#Total breakdowns. That is confirmed and unconfirmed breakdowns
model1T<-glm(Totalincidence~Treatment+Triplet+log(TotalHistoric)+log(Herds),poisson, data=RBCT)
model2T<-glm(Totalincidence~Triplet+log(TotalHistoric)+log(Herds),poisson, data=RBCT)
model3T<-glm(Totalincidence~Treatment+Triplet+log(TotalHistoric)+offset(log(Herds)),poisson, data=RBCT)
model4T<-glm(Totalincidence~Triplet+log(TotalHistoric)+offset(log(Herds)),poisson, data=RBCT)

unlist(lapply(list(model1T, model2T, model3T, model4T), AICc))
unlist(lapply(list(model1T, model2T, model3T, model4T), BICc))
aictab(list(model1T, model2T, model3T, model4T))

#Autocorrelation issues in all 4 models.
#Model1T was mention in the RBCT analysis, but a decision was made to stay with confirmed breakdowns only
check_autocorrelation(model1T)
check_autocorrelation(model2T)
check_autocorrelation(model3T)
check_autocorrelation(model4T)

#Moving to generalized poisson, with random effects and time at risk in the exposure variable
model1Tr<-glmmTMB(Totalincidence~Treatment+(1|Triplet)+log(TotalHistoric)+log(Herds*Timeatrisk),genpois, data=RBCT)
model2Tr<-glmmTMB(Totalincidence~(1|Triplet)+log(TotalHistoric)+log(Herds*Timeatrisk),genpois, data=RBCT)
model3Tr<-glmmTMB(Totalincidence~Treatment+(1|Triplet)+log(TotalHistoric)+offset(log(Herds*Timeatrisk)),genpois, data=RBCT)
model4Tr<-glmmTMB(Totalincidence~(1|Triplet)+log(TotalHistoric)+offset(log(Herds*Timeatrisk)),genpois, data=RBCT)

#Model2Tr is the most parsimonious
#Model 1TR does not have a significant  treatment effect
#beta for exposure is c 0.5
unlist(lapply(list(model1Tr, model2Tr, model3Tr, model4Tr), AICc))
unlist(lapply(list(model1Tr, model2Tr, model3Tr, model4Tr), BICc))
aictab(list(model1Tr, model2Tr, model3Tr, model4Tr))

#Most parsimonious model including Treatment co-variate, fails to reach the 5% significance
#for a treatment effect

summary(model1Tr)

#No issues with residuals
check_autocorrelation(model1Tr)
check_autocorrelation(model2Tr)
check_autocorrelation(model3Tr)
check_autocorrelation(model4Tr)

plot(simulateResiduals(model2Tr))

#Correcting for false positive unconfirmed breakdowns
#model1Tr:
# Number of simulations
n_sims <- 1000

# Matrix/dataframe to store the Treatment coefficient and p-value for each run
sim_results <- data.frame(
  Estimate = numeric(n_sims),
  StdError = numeric(n_sims),
  PValue   = numeric(n_sims)
)

set.seed(42) # For reproducibility

for(i in 1:n_sims) {
  # Copy dataframe
  temp_data <- RBCT
  
  # Sample 2 rows weighted by incidence
  fp_rows <- sample(1:nrow(temp_data), size = 2, replace = TRUE, prob = temp_data$Totalincidence)
  
  # Subtract the false positives
  for (r in fp_rows) {
    temp_data$Totalincidence[r] <- temp_data$Totalincidence[r] - 1
  }
  
  # Fit the model quietly
  mod <- tryCatch(
    glmmTMB(Totalincidence ~ Treatment + (1|Triplet) + log(TotalHistoric) + log(Herds*Timeatrisk), 
            family = genpois, data = temp_data),
    error = function(e) NULL
  )
  
  if(!is.null(mod)) {
    # 1. Grab the full conditional fixed-effects coefficient matrix
    coef_matrix <- summary(mod)$coefficients$cond
    
    # 2. Dynamically target the 2nd row (your Treatment effect) by index
    # This prevents any "subscript out of bounds" text-mismatch errors
    sim_results$Estimate[i] <- coef_matrix[2, "Estimate"]
    sim_results$StdError[i] <- coef_matrix[2, "Std. Error"]
    sim_results$PValue[i]   <- coef_matrix[2, "Pr(>|z|)"]
  }
}

# View the distribution of your Treatment effect across all 1,000 randomisations
summary(sim_results)

# Neighboring or perimeter zone: the perturbation effect.
#Model used to analyse the data in Donnelly et al 2007, and claimed a perturbation effect.
modelP<-glm(Neighbour_incidence~Treatment+Triplet+log(Neighbour_Historic)+log(Neighbour_Herds),poisson, data=RBCT)
summary(modelP)

# Modeled as generalised poisson, triplet as random effect and including time at risk in the exposure variable
#Competing models include the null model (no treatment) and offset models.
modelP1<-glmmTMB(Neighbour_incidence~Treatment+(1|Triplet)+log(Neighbour_Historic)+log(Neighbour_Herds*Timeatrisk),genpois, data=RBCT)
modelP2<-glmmTMB(Neighbour_incidence~(1|Triplet)+log(Neighbour_Historic)+log(Neighbour_Herds*Timeatrisk),genpois, data=RBCT)
modelP3<-glmmTMB(Neighbour_incidence~Treatment+(1|Triplet)+log(Neighbour_Historic)+offset(log(Neighbour_Herds*Timeatrisk)),genpois, data=RBCT)
modelP4<-glmmTMB(Neighbour_incidence~(1|Triplet)+log(Neighbour_Historic)+offset(log(Neighbour_Herds*Timeatrisk)),genpois, data=RBCT)

#modelP1 and modelP2 tied in terms of AICc. modelP2 has lowest BICc
unlist(lapply(list(modelP1, modelP2, modelP3, modelP4), AICc))
unlist(lapply(list(modelP1, modelP2, modelP3, modelP4), BICc))
aictab(list(modelP1, modelP2, modelP3, modelP4))

#modelP1 has issues with quantile deviations of residuals
check_autocorrelation(modelP1)
check_autocorrelation(modelP2)
 
plot(simulateResiduals(modelP1))
plot(simulateResiduals(modelP2))

#These models are further examined in the permutation models.
#See later

#Total breakdowns in surrounding areas as per RBCT histioric analysis
modelPT<-glm(Neighbour_Total_Incidence~Treatment+Triplet+log(Neighbour_Total_Historic)+log(Neighbour_Herds),poisson, data=RBCT)

#No effect of culling at 5% level. 
summary(modelPT)

#But issues with residuals
check_autocorrelation(modelPT)
plot(simulateResiduals(modelPT))

#Generalized poisson, random effects and time at risk in exposure variable
modelPT1<-glmmTMB(Neighbour_Total_Incidence~Treatment+(1|Triplet)+log(Neighbour_Total_Historic)+log(Neighbour_Herds),genpois, data=RBCT)
modelPT2<-glmmTMB(Neighbour_Total_Incidence~(1|Triplet)+log(Neighbour_Total_Historic)+log(Neighbour_Herds),genpois, data=RBCT)

#ModelPT2 has the lowest AICc and BICc. Neither model has an effect of treatment at 5%.
unlist(lapply(list(modelPT1, modelPT2), AICc))
unlist(lapply(list(modelPT1, modelPT2), BICc))
aictab(list(modelPT1, modelPT2))

#logistic regression models.
#Godfray et al analyses 4 models by logistic regression to conclude the RBCT results were sound.

RBCT$Historic.logodds = log(RBCT$Historic/(RBCT$Herds-RBCT$Historic))

modelB1 = glm(cbind(Incidence, Herds-Incidence)~ Treatment + Triplet + Historic.logodds,
              family=binomial, data=RBCT)
modelB2 = glm(cbind(Incidence, Herds-Incidence)~ Triplet + Historic.logodds, family=binomial,
              data=RBCT)
modelB3 = glm(cbind(Incidence, Herds-Incidence)~ Treatment + Triplet, offset=Historic.logodds,
              family=binomial, data=RBCT)
modelB4 = glm(cbind(Incidence, Herds-Incidence)~ Triplet, offset=Historic.logodds,
              family=binomial, data=RBCT)
unlist(lapply(list(modelB1, modelB2, modelB3, modelB4), AICc))


#Godfrey preferred model. Following selection by AICc, it was transformed to a quasipoisson for inference
modelB3q=glm(cbind(Incidence, Herds-Incidence)~ Treatment + Triplet, offset=Historic.logodds,
                 family=quasibinomial, data=RBCT)

summary(modelB3q)

#but modelB3 is NOT the most parsimonious by QAICc, and therefore invalidates simply transforming it to
#a quasipoisson for inference.
unlist(lapply(list(modelB1, modelB2, modelB3, modelB4), AICc, c.hat=modelB1$deviance/modelB1$df.resid))

#To increase the degrees of freedom, whilst retaining the block structure analysis
#Triplet as a random effect.
#To accommodate the time at risk, a cloglog link is introduced

RBCT$histcl<-log(-log(1-RBCT$Historic/RBCT$Herds))

modelB1r = glmmTMB(cbind(Incidence, Herds-Incidence)~ Treatment + (1|Triplet) +log(Timeatrisk)+histcl,
                   family=binomial(link="cloglog"), data=RBCT)
modelB2r = glmmTMB(cbind(Incidence, Herds-Incidence)~ (1|Triplet)+log(Timeatrisk)+histcl,
                   family=binomial(link="cloglog"), data=RBCT)
modelB3r = glmmTMB(cbind(Incidence, Herds-Incidence)~ Treatment + (1|Triplet) +log(Timeatrisk), offset=histcl,
              family=binomial(link="cloglog"), data=RBCT)
modelB4r = glmmTMB(cbind(Incidence, Herds-Incidence)~ (1|Triplet)+log(Timeatrisk), offset=histcl,
              family=binomial(link="cloglog"), data=RBCT)

#ModelB3r is the most parsimonious
unlist(lapply(list(modelB1r, modelB2r,modelB3r, modelB4r), AICc))
unlist(lapply(list(modelB1r, modelB2r,modelB3r, modelB4r), BICc))

#Substantial problems with residuals with modelB3r
check_autocorrelation(modelB3r)
plot(simulateResiduals(modelB3r))

#betabinomial family to control for possible over dispersion
modelB1ri = glmmTMB(cbind(Incidence, Herds-Incidence)~ Treatment + (1|Triplet) +log(Timeatrisk)+histcl,
             family=betabinomial(link="cloglog"), data=RBCT)
modelB2ri = glmmTMB(cbind(Incidence, Herds-Incidence)~ (1|Triplet)+log(Timeatrisk)+histcl,
             family=betabinomial(link="cloglog"), data=RBCT)
modelB3ri = glmmTMB(cbind(Incidence, Herds-Incidence)~ Treatment + (1|Triplet) +log(Timeatrisk), offset=histcl,
             family=betabinomial(link="cloglog"), data=RBCT)
modelB4ri = glmmTMB(cbind(Incidence, Herds-Incidence)~ (1|Triplet)+log(Timeatrisk), offset=histcl,
             family=betabinomial(link="cloglog"), data=RBCT)

#Little differences in AICc, although modelB3ri has the lowest
#Likewise with BICc, with model modelB4ri the lowest
unlist(lapply(list(modelB1ri, modelB2ri,modelB3ri, modelB4ri), AICc))
unlist(lapply(list(modelB1ri, modelB2ri,modelB3ri, modelB4ri), BICc))

check_autocorrelation(modelB1ri)
check_autocorrelation(modelB1ri)
check_autocorrelation(modelB1ri)
check_autocorrelation(modelB1ri)

#ModelB3ri and modelB43i have issues with quantile deviations
plot(simulateResiduals(modelB3ri))
plot(simulateResiduals(modelB4ri))

#A treatment effect is indicated with some of the binomial models, but not others that have similar AICc
#or BICc. There are issues with residuals in the most parsimonious of the models with a treatment effect
#All of the binomial models are invalid because the incidence was not a series of Bernoulli trials because of 
#Sequential multiple breakdowns in the same herd.

#When including total breakdowns, treatment effect disappears
RBCT$histclT<-log(-log(1-RBCT$TotalHistoric/RBCT$Herds))

modelB3riT = glmmTMB(cbind(Totalincidence, Herds-Totalincidence)~ Treatment + (1|Triplet) +log(Timeatrisk), offset=histclT,
                    family=betabinomial(link="cloglog"), data=RBCT)
modelB4riT = glmmTMB(cbind(Totalincidence, Herds-Totalincidence)~ (1|Triplet)+log(Timeatrisk), offset=histclT,
                    family=betabinomial(link="cloglog"), data=RBCT)

unlist(lapply(list(modelB3riT, modelB4riT), AICc))
unlist(lapply(list(modelB3riT, modelB4riT), BICc))
summary(modelB3riT)
summary(modelB4riT)

#However, both models are invalid as these are not Bernoulli trials
#Although modelB3riT has no issues with residuals.
check_autocorrelation(modelB3riT)
check_autocorrelation(modelB4riT)
plot(simulateResiduals(modelB3riT))
plot(simulateResiduals(modelB4riT))


# ==============================================================================
# PLoS Computation Biology - CORE PERMUTATION ENGINES
# Complete Implementation of Model A (Estimate) and Model B (Studentized Z)
# This is to analyse competing Poisson models (generalized poisson) for the 
# effect of Treatment on the incidence of herd breakdowns due to bTB.
# ==============================================================================

#Confirmed breakdowns (incidence) only, that is model1r
#Total breakdowns (incidence), that is model1Tr
#See main text for definitions of confirmed and total breakdowns

local({
  library(glmmTMB)
  library(dplyr)
  library(MuMIn)
  library(foreach)
  library(doParallel)
  library(doRNG)
  
  # FIXED: Removed comma separator so R parses the number as a valid integer
  nsim <- 10000
  
  # Exact column mapping matching primary model specifications
  scenarios <- list(
    Confirmed = list(target = "Incidence",      historic = "Historic"), 
    Total     = list(target = "Totalincidence", historic = "TotalHistoric") 
  )
  
  # Custom BICc function based on Ventura et al. (2019)
  BICc <- function(model) {
    LL <- logLik(object = model)
    df <- attr(LL, "df")
    n  <- length(residuals(model))
    return(as.numeric(-2 * LL + (n * log(n) * df) / (n - df - 2)))
  }
  
  # Container for side-by-side manuscript compilation
  final_results_table <- data.frame()
  
  # ==============================================================================
  # PARALLEL HARDWARE CONFIGURATION
  # ==============================================================================
  num_cores <- max(1, parallel::detectCores() - 1)
  cl <- makeCluster(num_cores)
  registerDoParallel(cl)
  
  # ==============================================================================
  # EXECUTION ENGINE
  # ==============================================================================
  
  for (scen_name in names(scenarios)) {
    cat("\n==================================================================\n")
    cat("RUNNING PARALLEL PERMUTATIONS FOR DATASET:", toupper(scen_name), "\n")
    cat("==================================================================\n")
    
    target_var   <- scenarios[[scen_name]]$target
    historic_var <- scenarios[[scen_name]]$historic
    
    # Dynamic formula formulation
    formula_full <- as.formula(paste0(target_var, " ~ Treatment + (1|Triplet) + log(", historic_var, ") + log(Herds*Timeatrisk)"))
    formula_null <- as.formula(paste0(target_var, " ~ (1|Triplet) + log(", historic_var, ") + log(Herds*Timeatrisk)"))
    
    # Fit Observed Models to derive baseline true targets
    model_full <- glmmTMB(formula_full, family = genpois, data = RBCT)
    model_null <- glmmTMB(formula_null, family = genpois, data = RBCT)
    
    obs_aicc_full <- MuMIn::AICc(model_full)
    obs_aicc_null <- MuMIn::AICc(model_null)
    obs_bicc_full <- BICc(model_full)
    obs_bicc_null <- BICc(model_null)
    
    all_coef_names <- names(fixef(model_full)$cond)
    treatment_key  <- all_coef_names[grep("Treatment", all_coef_names)]
    
    obs_estimate <- as.numeric(fixef(model_full)$cond[treatment_key])
    obs_z_val    <- as.numeric(coef(summary(model_full))$cond[treatment_key, "z value"])
    
    cat("Processing 10,000 iterations via distributed safe RNG stream...\n")
    
    # Set the reproducible seed right before the parallel step execution
    set.seed(42)
    
    # Parallel loop harvesting results into an isolated list structure
    perm_outputs <- foreach(
      i = 1:nsim,
      .packages = c("glmmTMB", "dplyr"),
      .export   = c("RBCT", "formula_full", "treatment_key")
    ) %dorng% {
      
      # Constrained Cluster Reshuffling: Shuffle Treatment only within Triplet boundaries
      permuted_data <- RBCT %>%
        group_by(Triplet) %>%
        mutate(Treatment = sample(as.character(Treatment))) %>%
        ungroup() %>%
        as.data.frame()
      
      # Fit the full model under the permuted data structure
      fit_perm <- try(
        glmmTMB(formula_full, family = genpois, data = permuted_data),
        silent = TRUE
      )
      
      # Initialize empty extraction structure
      out <- list(est = NA_real_, z = NA_real_, err_type = "success")
      
      if (inherits(fit_perm, "try-error")) {
        out$err_type <- "hard_error"
      } else if (fit_perm$fit$convergence != 0) {
        out$err_type <- "conv_failure"
      } else {
        coefs_matrix <- coef(summary(fit_perm))$cond
        if (treatment_key %in% rownames(coefs_matrix)) {
          out$est <- as.numeric(fixef(fit_perm)$cond[treatment_key])
          out$z   <- as.numeric(coefs_matrix[treatment_key, "z value"])
        }
      }
      out
    }
    
    # ==============================================================================
    # POST-PROCESSING HARVESTED METRICS
    # ==============================================================================
    perm_estimates <- sapply(perm_outputs, `[[`, "est")
    perm_z_values  <- sapply(perm_outputs, `[[`, "z")
    err_status     <- sapply(perm_outputs, `[[`, "err_type")
    
    hard_errors          <- sum(err_status == "hard_error")
    convergence_failures <- sum(err_status == "conv_failure")
    
    valid_indices   <- !is.na(perm_estimates)
    clean_estimates <- perm_estimates[valid_indices]
    clean_z_values  <- perm_z_values[valid_indices]
    n_success       <- length(clean_estimates)
    
    # LOCALIZED DIAGNOSTIC HEALTH REPORT
    cat("\n--- ITERATION REPORT (", scen_name, ") ---\n")
    cat("  Total Successful Clean Cycles :", n_success, "out of", nsim, "\n")
    cat("  Kicked-out Code Crashes       :", hard_errors, "\n")
    cat("  Kicked-out Non-Convergence   :", convergence_failures, "\n")
    
    # Calculate Two-Sided Empirical Probabilities
    p_val_model_a <- mean(abs(clean_estimates) >= abs(obs_estimate)) # Parameter Path
    p_val_model_b <- mean(abs(clean_z_values) >= abs(obs_z_val))     # Wald z Path
    
    # Derive Clopper-Pearson 95% Confidence Intervals
    ci_a <- binom.test(sum(abs(clean_estimates) >= abs(obs_estimate)), n_success)$conf.int
    ci_b <- binom.test(sum(abs(clean_z_values) >= abs(obs_z_val)), n_success)$conf.int
    
    # FIXED: Corrected indices to print [Lower Bound (1), Upper Bound (2)] accurately
    scen_summary <- data.frame(
      Dataset               = scen_name,
      Observed_IRR          = round(exp(obs_estimate), 4),
      Observed_Z            = round(obs_z_val, 3),
      Delta_AICc            = round(obs_aicc_full - obs_aicc_null, 2),
      Delta_BICc            = round(obs_bicc_full - obs_bicc_null, 2),
      Model_A_P             = round(p_val_model_a, 4),
      Model_A_CI            = paste0("[", round(ci_a[1], 4), ", ", round(ci_a[2], 4), "]"),
      Model_B_P             = round(p_val_model_b, 4),
      Model_B_CI            = paste0("[", round(ci_b[1], 4), ", ", round(ci_b[2], 4), "]")
    )
    
    final_results_table <- rbind(final_results_table, scen_summary)
  }
  
  # Close down hardware nodes cleanly
  stopCluster(cl)
  
  # ==============================================================================
  # DISPLAY INTEGRATED MATRIX RESULTS
  # ==============================================================================
  cat("\n==================================================================\n")
  cat("COMPLETED SIDE-BY-SIDE ANALYTICAL GRID (10,000 PERMUTATIONS):\n")
  cat("==================================================================\n")
  print(final_results_table, row.names = FALSE)
})

  # ==============================================================================
  #INCIDENCE IN NEIGHBORING AREAS: THE "PERTURBATION" EFFECT
  #Confirmed breakdowns (incidence) only, that is modelP1
  #and models with total breakdowns - modelP1, and with offsets
  #Including full screening of competing models
  # ==============================================================================

local({
  library(glmmTMB)
  library(dplyr)
  library(MuMIn)
  library(foreach)
  library(doParallel)
  library(doRNG)
  
  # Ensure Triplet is treated as a clean categorical factor
  RBCT$Triplet <- as.factor(RBCT$Triplet)
  
  # Custom BICc function based on Ventura et al. (2019)
  BICc <- function(model) {
    LL <- logLik(object = model)
    df <- attr(LL, "df")
    n  <- length(residuals(model))
    return(as.numeric(-2 * LL + (n * log(n) * df) / (n - df - 2)))
  }
  
  # Simulation Parameters - Scaled to 10,000 for maximum statistical power
  nsim <- 10000
  
  # EXACT NEIGHBOUR PERIMETER SCENARIO DICTIONARY
  scenarios_P <- list(
    Confirmed_Neighbour = list(target = "Neighbour_incidence",       historic = "Neighbour_Historic"), 
    Total_Neighbour     = list(target = "Neighbour_Total_Incidence", historic = "Neighbour_Total_Historic") 
  )
  
  # Container for side-by-side manuscript compilation
  perturbation_results_table <- data.frame()
  
  # ==============================================================================
  # PARALLEL HARDWARE CONFIGURATION
  # ==============================================================================
  num_cores <- max(1, parallel::detectCores() - 1)
  cl <- makeCluster(num_cores)
  registerDoParallel(cl)
  
  # ==============================================================================
  # EXECUTION ENGINE
  # ==============================================================================
  
  for (scen_name in names(scenarios_P)) {
    cat("\n==================================================================\n")
    cat("PROCESSING PERIMETER DATASET:", toupper(scen_name), "\n")
    cat("==================================================================\n")
    
    target_var   <- scenarios_P[[scen_name]]$target
    historic_var <- scenarios_P[[scen_name]]$historic
    
    # 1. PRE-SCREEN EXPOSURE SCALING
    mP1 <- glmmTMB(as.formula(paste0(target_var, " ~ Treatment + (1|Triplet) + log(", historic_var, ") + log(Neighbour_Herds*Timeatrisk)")), family = genpois, data = RBCT)
    mP2 <- glmmTMB(as.formula(paste0(target_var, " ~ (1|Triplet) + log(", historic_var, ") + log(Neighbour_Herds*Timeatrisk)")), family = genpois, data = RBCT)
    mP3 <- glmmTMB(as.formula(paste0(target_var, " ~ Treatment + (1|Triplet) + log(", historic_var, ") + offset(log(Neighbour_Herds*Timeatrisk))")), family = genpois, data = RBCT)
    mP4 <- glmmTMB(as.formula(paste0(target_var, " ~ (1|Triplet) + log(", historic_var, ") + offset(log(Neighbour_Herds*Timeatrisk))")), family = genpois, data = RBCT)
    
    aicc_vector <- unlist(lapply(list(mP1, mP2, mP3, mP4), AICc))
    cat("AICc Model Prescreen Profile:\n")
    cat("  P1 (Full, Explanatory Exposure Slope) :", round(aicc_vector[1], 2), "\n")
    cat("  P2 (Null, Explanatory Exposure Slope) :", round(aicc_vector[2], 2), "\n")
    cat("  P3 (Full, Rigid Fixed Exposure Offset) :", round(aicc_vector[3], 2), "\n")
    cat("  P4 (Null, Rigid Fixed Exposure Offset) :", round(aicc_vector[4], 2), "\n")
    
    # Automated routing
    if (min(aicc_vector[1], aicc_vector[2]) < min(aicc_vector[3], aicc_vector[4])) {
      cat("--> Selection Analysis: Explanatory Exposure Framework is preferred (Saturated).\n")
      formula_full <- as.formula(paste0(target_var, " ~ Treatment + (1|Triplet) + log(", historic_var, ") + log(Neighbour_Herds*Timeatrisk)"))
      model_full   <- mP1
      model_null   <- mP2
    } else {
      cat("--> Selection Analysis: Fixed Offset Framework is preferred (Linear).\n")
      formula_full <- as.formula(paste0(target_var, " ~ Treatment + (1|Triplet) + log(", historic_var, ") + offset(log(Neighbour_Herds*Timeatrisk))"))
      model_full   <- mP3
      model_null   <- mP4
    }
    
    # Extract true observed targets
    obs_aicc_full <- AICc(model_full)
    obs_aicc_null <- AICc(model_null)
    obs_bicc_full <- BICc(model_full)
    obs_bicc_null <- BICc(model_null)
    
    all_coef_names <- names(fixef(model_full)$cond)
    treatment_key  <- all_coef_names[grep("Treatment", all_coef_names)]
    
    obs_estimate <- as.numeric(fixef(model_full)$cond[treatment_key])
    obs_z_val    <- as.numeric(coef(summary(model_full))$cond[treatment_key, "z value"])
    
    cat("Processing", nsim, "parallel permutations via distributed safe RNG stream...\n")
    
    # Synchronize the seed directly prior to parallel collection
    set.seed(42)
    
    # Parallelize while allowing doRNG to handle explicit workspace variables
    perm_outputs <- foreach(
      i = 1:nsim,
      .packages = c("glmmTMB", "dplyr")
    ) %dorng% {
      
      # Block Constraints: Reshuffle treatment assignments exclusively within Triplet blocks
      permuted_data <- RBCT %>%
        group_by(Triplet) %>%
        mutate(Treatment = sample(as.character(Treatment))) %>%
        ungroup() %>%
        as.data.frame()
      
      # Fit model structure to reshuffled data partition
      fit_perm <- try(
        glmmTMB(formula_full, family = genpois, data = permuted_data),
        silent = TRUE
      )
      
      out <- list(est = NA_real_, z = NA_real_, err_type = "success")
      
      if (inherits(fit_perm, "try-error")) {
        out$err_type <- "hard_error"
      } else if (fit_perm$fit$convergence != 0) {
        out$err_type <- "conv_failure"
      } else {
        coefs_matrix <- coef(summary(fit_perm))$cond
        if (treatment_key %in% rownames(coefs_matrix)) {
          out$est <- as.numeric(fixef(fit_perm)$cond[treatment_key])
          out$z   <- as.numeric(coefs_matrix[treatment_key, "z value"])
        }
      }
      out
    }
    
    # ==============================================================================
    # POST-PROCESSING HARVESTED METRICS
    # ==============================================================================
    perm_estimates <- sapply(perm_outputs, `[[`, "est")
    perm_z_values  <- sapply(perm_outputs, `[[`, "z")
    err_status     <- sapply(perm_outputs, `[[`, "err_type")
    
    hard_errors          <- sum(err_status == "hard_error")
    convergence_failures <- sum(err_status == "conv_failure")
    
    valid_indices   <- !is.na(perm_estimates)
    clean_estimates <- perm_estimates[valid_indices]
    clean_z_values  <- perm_z_values[valid_indices]
    n_success       <- length(clean_estimates)
    
    cat("\n--- PERIMETER HEALTH REPORT (", scen_name, ") ---\n")
    cat("  Total Successful Clean Cycles :", n_success, "out of", nsim, "\n")
    cat("  Kicked-out Code Crashes       :", hard_errors, "\n")
    cat("  Kicked-out Non-Convergence   :", convergence_failures, "\n")
    
    # Calculate Two-Sided Empirical Probabilities
    p_val_model_a <- mean(abs(clean_estimates) >= abs(obs_estimate))
    p_val_model_b <- mean(abs(clean_z_values) >= abs(obs_z_val))
    
    # Compute Exact Clopper-Pearson 95% Confidence Intervals
    ci_a <- binom.test(sum(abs(clean_estimates) >= abs(obs_estimate)), n_success)$conf.int
    ci_b <- binom.test(sum(abs(clean_z_values) >= abs(obs_z_val)), n_success)$conf.int
    
    # Compile metrics into Model A and Model B labeling paths
    scen_summary <- data.frame(
      Dataset      = scen_name,
      Observed_IRR = round(exp(obs_estimate), 4),
      Observed_Z   = round(obs_z_val, 3),
      Delta_AICc   = round(obs_aicc_full - obs_aicc_null, 2),
      Delta_BICc   = round(obs_bicc_full - obs_bicc_null, 2),
      Model_A_P    = round(p_val_model_a, 4),
      Model_A_CI   = paste0("[", round(ci_a[1], 4), ", ", round(ci_a[2], 4), "]"),
      Model_B_P    = round(p_val_model_b, 4),
      Model_B_CI   = paste0("[", round(ci_b[1], 4), ", ", round(ci_b[2], 4), "]")
    )
    
    perturbation_results_table <- rbind(perturbation_results_table, scen_summary)
  }
  
  # Always dismantle backend nodes smoothly
  stopCluster(cl)
  
  # ==============================================================================
  # DISPLAY INTEGRATED PERIMETER MATRIX RESULTS
  # ==============================================================================
  cat("\n==================================================================\n")
  cat("COMPLETED SIDE-BY-SIDE PERIMETER ANALYTICAL GRID (10,000 SIMULATIONS):\n")
  cat("==================================================================\n")
  print(perturbation_results_table, row.names = FALSE)
})


# ==============================================================================
# EXPERIMENTAL DESIGN & RESAMPLING DIAGNOSTICS: CLUSTER BLOCK BOOTSTRAP
# Verification of Model Parsimony and Post-Processing Leverage Engines
# ==============================================================================


library(glmmTMB)
library(dplyr)
library(MuMIn)
library(ggplot2)
library(foreach)
library(doParallel)
library(doRNG)

# Ensure Triplet factor vector is set
RBCT$Triplet <- as.factor(RBCT$Triplet)
triplet_ids  <- unique(RBCT$Triplet)

# Simulation Parameters
nsim <- 10000
set.seed(42)


# ==============================================================================
# PARALLEL CONFIGURATION to increase speed
# ==============================================================================
# Use 1 less than total cores to prevent system freezing
num_cores <- max(1, parallel::detectCores() - 1)
cl <- makeCluster(num_cores)
registerDoParallel(cl)

cat("Running 10,000 block-resampled models in parallel across", num_cores, "cores...\n")

# Use %dopar% for parallel execution. Ensure required packages are exported.
parallel_results <- foreach(
  i = 1:nsim, 
  .packages = c("glmmTMB", "dplyr", "MuMIn"),
  .export = c("triplet_ids", "RBCT")
) %dorng% {
  
  # Sample Triplet blocks with replacement
  sampled_triplets <- data.frame(Triplet = sample(triplet_ids, replace = TRUE))
  
  # Log frequency table for this specific iteration
  counts_vector <- table(factor(sampled_triplets$Triplet, levels = triplet_ids))
  
  # Reconstruct dataset mapping separate structural identifiers
  boot_data <- sampled_triplets %>%
    mutate(New_Cluster = row_number()) %>%
    left_join(RBCT, by = "Triplet", relationship = "many-to-many") %>%
    as.data.frame()
  
  # Fit Model 1 (Full) vs Model 2 (Null)
  fit1 <- try(glmmTMB(Incidence ~ Treatment + (1|New_Cluster) + log(Historic) + log(Herds*Timeatrisk), family = genpois, data = boot_data), silent = TRUE)
  fit2 <- try(glmmTMB(Incidence ~ (1|New_Cluster) + log(Historic) + log(Herds*Timeatrisk), family = genpois, data = boot_data), silent = TRUE)
  
  # Default empty result line
  metrics <- c(AIC_Full=NA, AIC_Null=NA, Delta_AIC=NA, AICc_Full=NA, AICc_Null=NA, Delta_AICc=NA)
  
  if (!inherits(fit1, "try-error") && !inherits(fit2, "try-error")) {
    aic_1  <- AIC(fit1)
    aic_2  <- AIC(fit2)
    aicc_1 <- MuMIn::AICc(fit1)
    aicc_2 <- MuMIn::AICc(fit2)
    
    metrics <- c(aic_1, aic_2, (aic_1 - aic_2), aicc_1, aicc_2, (aicc_1 - aicc_2))
  }
  
  # Return both things as a list for this iteration
  list(metrics = metrics, counts = counts_vector)
}

# ALWAYS shut down the parallel cluster when finished
stopCluster(cl)

# ==============================================================================
# POST-PROCESSING
# ==============================================================================
# Extract metrics into a clean data frame
boot_aic_results <- do.call(rbind, lapply(parallel_results, `[[`, "metrics"))
colnames(boot_aic_results) <- c("AIC_Full", "AIC_Null", "Delta_AIC", "AICc_Full", "AICc_Null", "Delta_AICc")

boot_df <- as.data.frame(boot_aic_results)
boot_df$Iteration_Index <- 1:nsim
boot_df <- na.omit(boot_df)
n_valid <- nrow(boot_df)

# Extract counts into a frequency matrix
triplet_tracker <- lapply(parallel_results, `[[`, "counts")
all_triplets <- as.character(triplet_ids)

frequency_matrix <- do.call(rbind, lapply(triplet_tracker, function(counts) {
  vec <- rep(0, length(all_triplets))
  names(vec) <- all_triplets
  vec[names(counts)] <- as.numeric(counts)
  return(vec)
}))

# ==============================================================================
# DATA SUMMARY PRINTING
# ==============================================================================
pct_full_better_aicc <- mean(boot_df$Delta_AICc <= -2) * 100
pct_tied_aicc        <- mean(boot_df$Delta_AICc > -2 & boot_df$Delta_AICc < 2) * 100
pct_null_better_aicc <- mean(boot_df$Delta_AICc >= 2) * 100

cat("\n--- BOOTSTRAP MODEL SELECTION METRICS (N =", n_valid, ") ---\n\n")
cat("Full Model Wins (AICc <= -2)     :", round(pct_full_better_aicc, 1), "%\n")
cat("Models are Tied (-2 < AICc < 2)  :", round(pct_tied_aicc, 1), "%\n")
cat("Null Model Wins (AICc >= 2)      :", round(pct_null_better_aicc, 1), "%\n")

# ==============================================================================
# TAIL-LEVERAGE EVALUATION ENGINE
# ==============================================================================
tracking_matrix <- frequency_matrix[boot_df$Iteration_Index, , drop = FALSE]
tail_indices <- which(boot_df$Delta_AICc <= -2)

expected_appearance <- colMeans(tracking_matrix)
avg_tail_appearance <- colMeans(tracking_matrix[tail_indices, , drop = FALSE])

leverage_table <- data.frame(
  Triplet_ID              = triplet_ids,
  Avg_Appearance_In_Tail  = round(avg_tail_appearance, 2),
  Expected_Appearance     = round(expected_appearance, 2),
  Over_Representation_Ratio = round(avg_tail_appearance / expected_appearance, 2)
) %>% arrange(desc(Over_Representation_Ratio))

cat("\n--- TRACKED CLUSTER BLOCK LEVERAGE METRICS ---\n")
print(leverage_table, row.names = FALSE)

# ==============================================================================
# FINAL DIAGNOSTIC MATRIX FIX
# ==============================================================================
diagnostic_df <- as.data.frame(frequency_matrix)
diagnostic_df <- diagnostic_df[boot_df$Iteration_Index, , drop = FALSE]
diagnostic_df$Delta_AICc <- boot_df$Delta_AICc

# ==============================================================================
# CEILING-BALANCED GGPLOT2 PLOT ASSEMBLY
# ==============================================================================
max_density <- max(density(boot_df$Delta_AICc)$y)
label_y     <- max_density * 0.95

p <- ggplot(boot_df, aes(x = Delta_AICc)) +
  annotate("rect", xmin = -Inf, xmax = -2, ymin = 0, ymax = Inf, fill = "#228B22", alpha = 0.15) + 
  annotate("rect", xmin = -2, xmax = 2, ymin = 0, ymax = Inf, fill = "#FFE4E1", alpha = 0.3) +    
  annotate("rect", xmin = 2, xmax = Inf, ymin = 0, ymax = Inf, fill = "#00008B", alpha = 0.1) +   
  geom_histogram(aes(y = after_stat(density)), bins = 40, fill = "grey40", color = "white", alpha = 0.7) +
  geom_density(color = "black", linewidth = 1) +
  geom_vline(xintercept = 0, linetype = "solid", color = "red", linewidth = 0.8) +
  geom_vline(xintercept = c(-2, 2), linetype = "dashed", color = "grey30", linewidth = 0.6) +
  
  annotate("text", x = min(boot_df$Delta_AICc)*0.6, y = label_y, label = "Full Model Wins", color = "#1e5a1e", fontface = "bold", size = 3.5) +
  annotate("text", x = 0, y = label_y * 1.05, label = "Models Tied\n(Favors Null)", color = "brown", fontface = "bold", size = 3.5) +
  annotate("text", x = max(boot_df$Delta_AICc)*0.6, y = label_y, label = "Null Wins", color = "#00005b", fontface = "bold", size = 3.5) +
  
  labs(
    title = "Bootstrap Distribution of Delta AICc",
    subtitle = "Formula: AICc(Full Model) - AICc(Null Model)",
    x = expression(paste(Delta, "AICc (Negative values favor Full Model)")),
    y = "Density"
  ) +
  theme_minimal(base_size = 13) +
  theme(
    plot.title = element_text(face = "bold", hjust = 0.5), 
    plot.subtitle = element_text(color = "grey40", hjust = 0.5)
  )

print(p)

all_triplets <- as.character(triplet_ids)

# Vectorized conversion of list tables into a clean frequency matrix
frequency_matrix <- do.call(rbind, lapply(triplet_tracker, function(counts) {
  vec <- rep(0, length(all_triplets))
  names(vec) <- all_triplets
  vec[names(counts)] <- as.numeric(counts)
  return(vec)
}))

diagnostic_df <- as.data.frame(frequency_matrix)

# FIX: Map only the valid bootstrap results using the matching Iteration Index
diagnostic_df <- diagnostic_df[boot_df$Iteration_Index, , drop = FALSE]
diagnostic_df$Delta_AICc <- boot_df$Delta_AICc

# ==============================================================================
# 4. ISOLATE THE LEFT TAIL (Dynamic Quantile Inversion for JRSS Rigor)
# ==============================================================================
# Defines the tail objectively as the lowest 5% of all information swings
tail_quantile  <- 0.05 
tail_threshold <- as.numeric(quantile(diagnostic_df$Delta_AICc, probs = tail_quantile))

left_tail_iterations <- diagnostic_df %>% filter(Delta_AICc <= tail_threshold)

# ==============================================================================
# 5. CALCULATE THE AVERAGE MULTIPLICITY IN THE LEFT TAIL VS. OVERALL
# ==============================================================================
cat("--- OUTLIER CLUSTER DIAGNOSTIC REPORT ---\n")
cat("Empirical Left-Tail Cutoff Threshold (", tail_quantile * 100, "% Quantile) :", round(tail_threshold, 2), "\n")
cat("Number of bootstrap iterations falling into the extreme left tail   :", nrow(left_tail_iterations), "\n\n")

if(nrow(left_tail_iterations) == 0) {
  cat("Error: Tail partition space is empty. Check underlying model convergence patterns.\n")
} else {
  # Extract appearance vectors matching your exact calculations
  tail_profile     <- colMeans(left_tail_iterations[, all_triplets])
  baseline_profile <- colMeans(diagnostic_df[, all_triplets]) 
  importance_ratio <- tail_profile / baseline_profile
  
  results_table <- data.frame(
    Triplet_ID                = names(tail_profile),
    Avg_Appearance_In_Tail    = round(tail_profile, 2),
    Expected_Appearance       = round(baseline_profile, 2),
    Over_Representation_Ratio = round(importance_ratio, 2)
  ) %>% arrange(desc(Over_Representation_Ratio))
  
  # Highlight the severe unbalance of Triplets C and A for publication records
  print(results_table)
}

library(ggplot2)
library(dplyr)
library(patchwork)
local({
  library(ggplot2)
  library(dplyr)
  library(patchwork)
  
  orr_df <- data.frame(
    ID  = c("C", "A", "F", "J", "E", "I", "G", "B", "D", "H"),
    Val = c(1.59, 1.44, 1.38, 1.32, 1.11, 1.11, 1.05, 0.76, 0.12, 0.07)
  )
  
  orr_df <- orr_df %>%
    mutate(
      ID = factor(ID, levels = rev(c("C", "A", "F", "J", "E", "I", "G", "B", "D", "H"))),
      Tier = case_when(
        Val >= 1.25 ~ "Core Driver",
        Val >= 0.90 ~ "Neutral Background",
        TRUE        ~ "Structural Blocker"
      ),
      Tier = factor(Tier, levels = c("Core Driver", "Neutral Background", "Structural Blocker"))
    )
  
  # ==============================================================================
  # PANEL A: DENSITY PATTERN WITH OVERLAP FIX
  # ==============================================================================
  panel_a <- ggplot(boot_df, aes(x = Delta_AICc)) +
    annotate("rect", xmin = -Inf, xmax = -2, ymin = 0, ymax = Inf, fill = "#228B22", alpha = 0.12) + 
    annotate("rect", xmin = -2, xmax = 2, ymin = 0, ymax = Inf, fill = "#FFE4E1", alpha = 0.25) +    
    annotate("rect", xmin = 2, xmax = Inf, ymin = 0, ymax = Inf, fill = "#00008B", alpha = 0.08) +   
    geom_histogram(aes(y = after_stat(density)), bins = 50, fill = "grey40", color = "white", alpha = 0.6) +
    geom_density(color = "black", linewidth = 0.9) +
    geom_vline(xintercept = 0, linetype = "solid", color = "red", linewidth = 0.8) +
    geom_vline(xintercept = c(-2, 2), linetype = "dashed", color = "grey30", linewidth = 0.6) +
    
    # FIXED: Using label instead of text with white backgrounds to eliminate collision
    annotate("label", x = -50, y = 0.083, label = "Full Model Wins", color = "#1e5a1e", fontface = "bold", size = 3.2, label.size = NA, fill = "white", alpha = 0.8) +
    annotate("label", x = 0, y = 0.083, label = "Models Tied\n(Favors Null)", color = "brown", fontface = "bold", size = 3.2, label.size = NA, fill = "white", alpha = 0.8) +
    annotate("label", x = 16, y = 0.083, label = "Null Wins", color = "#00005b", fontface = "bold", size = 3.2, label.size = NA, fill = "white", alpha = 0.8) +
    
    labs(
      title = "Bootstrap Distribution of Delta AICc (B = 10,000)",
      subtitle = "Formula: AICc(Full Model) - AICc(Null Model)",
      x = expression(paste(Delta, "AICc (Negative values favor Full Model)")),
      y = "Density"
    ) +
    theme_minimal(base_size = 12) +
    theme(plot.title = element_text(face = "bold", size = 13), axis.title.x = element_blank())
  
  # PANEL B: LEVERAGE BARS
  panel_b <- ggplot(orr_df, aes(x = ID, y = Val, fill = Tier)) +
    geom_bar(stat = "identity", width = 0.75, color = "black", linewidth = 0.3) +
    geom_hline(yintercept = 1.0, linetype = "dashed", color = "red", linewidth = 0.8) +
    geom_text(aes(label = sprintf("%.2f", Val)), hjust = -0.2, fontface = "bold", size = 3.5) +
    coord_flip() +
    scale_fill_manual(values = c("Core Driver" = "#228B22", "Neutral Background" = "grey80", "Structural Blocker" = "#00008B")) +
    scale_y_continuous(limits = c(0, 1.85), breaks = seq(0, 1.5, by = 0.5)) +
    labs(
      title = "Cluster-Block Tail Over-Representation Metrics",
      subtitle = "Values above 1.0 drive the long left selection tail",
      x = "Triplet Identifier",
      y = "Over-Representation Ratio (ORR)",
      fill = "Cluster Structural Class"
    ) +
    theme_minimal(base_size = 12) +
    theme(plot.title = element_text(face = "bold", size = 13), legend.position = "bottom", panel.grid.minor = element_blank())
  
  out_fig <- panel_a / panel_b + plot_annotation(tag_levels = 'A') & theme(plot.tag = element_text(face = 'bold', size = 16))
  print(out_fig)
})

# ==============================================================================
# 1. GENERATE STATISTICAL SUMMARY
# ==============================================================================
# Target the extreme levers identified by your bootstrap analysis
target_triplets <- c("C", "A", "D", "H")

library(dplyr)
library(ggplot2)


cluster_summary <- RBCT %>%
  filter(Triplet %in% target_triplets) %>%
  group_by(Triplet, Treatment) %>%
  summarize(
    Total_Incidence = sum(Incidence, na.rm = TRUE),
    Total_Herds     = sum(Herds, na.rm = TRUE),
    Mean_Incidence  = mean(Incidence, na.rm = TRUE),
    Total_Time      = sum(Timeatrisk, na.rm = TRUE),
    Row_Count       = n(),
    .groups = "drop"
  )

cat("--- RAW DATA SUMMARY FOR EXTREME LEVERAGE CLUSTERS ---\n")
print(cluster_summary)

# ==============================================================================
# 2. ASSEMBLE COMPARISON PLOT
# ==============================================================================
# Filter dataset down to the key clusters for clean plotting
plot_data <- RBCT %>% 
  filter(Triplet %in% target_triplets)

p_leverage <- ggplot(plot_data, aes(x = Treatment, y = Incidence, fill = Treatment)) +
  # Use boxplots to check the distributions
  geom_boxplot(alpha = 0.6, outlier.shape = NA, color = "grey20") +
  # Overlay raw data points to reveal sample sizes and exact distribution shapes
  geom_jitter(width = 0.2, height = 0, alpha = 0.5, size = 2) +
  # Split the plot into separate panels per cluster
  facet_wrap(~ Triplet, scales = "free_y") +
  scale_fill_manual(values = c("Control" = "#E69F00", "Treatment" = "#56B4E9")) +
  labs(
    title = "Raw Data Dissection of High vs. Low Leverage Clusters",
    subtitle = "Top Row: Full Model Drivers (C, A) | Bottom Row: Full Model Killers (D, H)",
    x = "Experimental Group",
    y = "Incidence Counts",
    fill = "Group"
  ) +
  theme_bw(base_size = 14) +
  theme(
    plot.title = element_text(face = "bold", hjust = 0.5),
    plot.subtitle = element_text(color = "grey40", hjust = 0.5),
    strip.background = element_rect(fill = "grey95"),
    strip.text = element_text(face = "bold", size = 14),
    legend.position = "bottom"
  )

print(p_leverage)


library(glmmTMB)
library(dplyr)
library(MuMIn)

# Ensure types are correct
RBCT$Treatment <- as.factor(RBCT$Treatment)
RBCT$Triplet   <- as.factor(RBCT$Triplet)
all_triplets   <- unique(RBCT$Triplet)

# Baseline model metrics (with all data)
base_model <- glmmTMB(
  Incidence ~ Treatment + (1|Triplet) + log(Historic) + log(Herds*Timeatrisk), 
  family = genpois, 
  data = RBCT
)

sum_base  <- summary(base_model)$coefficients$cond
treat_row <- rownames(sum_base)[grep("Treatment", rownames(sum_base))]

# Initialize baseline row
loo_results <- data.frame(
  Dropped_Cluster = "NONE (Baseline)",
  Estimate        = round(sum_base[treat_row, "Estimate"], 4),
  Std_Error       = round(sum_base[treat_row, "Std. Error"], 4),
  Z_Value         = round(sum_base[treat_row, "z value"], 2),
  P_Value         = round(sum_base[treat_row, "Pr(>|z|)"], 4),
  AICc            = round(MuMIn::AICc(base_model), 1),
  Flipped_Signif  = "No"
)

# ==============================================================================
# EXECUTE LOO ITERATION LOOP
# ==============================================================================
for (cluster in sort(all_triplets)) {
  
  # Subset to drop exactly one cluster
  sub_data <- filter(RBCT, Triplet != cluster)
  
  # Fit the model on the remaining data
  fit_loo <- try(glmmTMB(
    Incidence ~ Treatment + (1|Triplet) + log(Historic) + log(Herds*Timeatrisk), 
    family = genpois, 
    data = sub_data
  ), silent = TRUE)
  
  if (!inherits(fit_loo, "try-error")) {
    sum_loo <- summary(fit_loo)$coefficients$cond
    p_val   <- sum_loo[treat_row, "Pr(>|z|)"]
    
    # Check if this dropped cluster flips the significance threshold (0.05)
    is_flipped <- if ((sum_base[treat_row, "Pr(>|z|)"] <= 0.05 && p_val > 0.05) || 
                      (sum_base[treat_row, "Pr(>|z|)"] > 0.05 && p_val <= 0.05)) "YES" else "No"
    
    # Append results
    loo_results <- rbind(loo_results, data.frame(
      Dropped_Cluster = paste("Dropped", cluster),
      Estimate        = round(sum_loo[treat_row, "Estimate"], 4),
      Std_Error       = round(sum_loo[treat_row, "Std. Error"], 4),
      Z_Value         = round(sum_loo[treat_row, "z value"], 2),
      P_Value         = round(p_val, 4),
      AICc            = round(MuMIn::AICc(fit_loo), 1),
      Flipped_Signif  = is_flipped
    ))
  } else {
    # Log model optimization failures if any occur
    loo_results <- rbind(loo_results, data.frame(
      Dropped_Cluster = paste("Dropped", cluster),
      Estimate = NA, Std_Error = NA, Z_Value = NA, P_Value = NA, AICc = NA, Flipped_Signif = "Error"
    ))
  }
}

# ==============================================================================
# PRINT FINAL DIAGNOSTICS
# ==============================================================================
cat("\n--- LEAVE-ONE-OUT (LOO) SENSITIVITY MATRIX ---\n")
print(loo_results, row.names = FALSE)
```
